# The BNT162b2 mRNA SARS-CoV-2 vaccine induces transient afucosylated IgG1 in naive but not antigen-experienced vaccinees

**DOI:** 10.1101/2022.02.14.480353

**Authors:** Julie Van Coillie, Tamas Pongracz, Johann Rahmöller, Hung-Jen Chen, Chiara Geyer, Lonneke A. van Vlught, Jana S. Buhre, Tonći Šuštić, Thijs L. J. van Osch, Maurice Steenhuis, Willianne Hoepel, Wenjun Wang, Anne S. Lixenfeld, Jan Nouta, Sofie Keijzer, Federica Linty, Remco Visser, Mads D. Larsen, Emily L. Martin, Inga Künsting, Selina Lehrian, Vera von Kopylow, Carsten Kern, Hanna B. Lunding, Menno de Winther, Niels van Mourik, Theo Rispens, Tobias Graf, Marleen A. Slim, René Minnaar, Marije K. Bomers, Jonne J. Sikkens, Alexander P. J. Vlaar, C. Ellen van der Schoot, Jeroen den Dunnen, Manfred Wuhrer, Marc Ehlers, Gestur Vidarsson, the Fatebenefratelli-Sacco Infectious Diseases Physicians group and UMC COVID-19 S3/HCW study group

**Affiliations:** Department of Experimental Immunohematology, Sanquin Research, Amsterdam, The Netherlands; Landsteiner Laboratory, Amsterdam UMC, University of Amsterdam, Amsterdam, The Netherlands; Center for Proteomics and Metabolomics, Leiden University Medical Center, Leiden, The Netherlands; Laboratories of Immunology and Antibody Glycan Analysis, Institute of Nutritional Medicine, University of Lübeck and University Medical Center of Schleswig-Holstein, Lübeck, Germany; Department of Anesthesiology and Intensive Care, University of Lübeck and University Medical Center of Schleswig-Holstein, Lübeck, Germany; Center for Experimental and Molecular Medicine, Amsterdam Infection & Immunity Institute, Amsterdam, The Netherlands; Department of Medical Biochemistry, Experimental Vascular Biology, Amsterdam Cardiovascular Sciences, Amsterdam Infection and Immunity, Amsterdam UMC, University of Amsterdam, The Netherlands; Department of Intensive Care, Amsterdam University Medical Centers, University of Amsterdam, Amsterdam, The Netherlands; Department of Immunopathology, Sanquin Research, Amsterdam, The Netherlands; Department of Experimental Immunology, Amsterdam UMC, University of Amsterdam, Amsterdam, The Netherlands; Department of Rheumatology and Clinical Immunology, Amsterdam UMC, Amsterdam Rheumatology and Immunology Center, Amsterdam, The Netherlands; Medical Department 3, University Medical Center of Schleswig-Holstein, Lübeck, Germany; Amsterdam UMC Biobank, Amsterdam UMC, Amsterdam, The Netherlands; Department of Internal Medicine, Amsterdam Infection and Immunity Institute, Amsterdam UMC, Vrije Universiteit Amsterdam; Airway Research Center North, University of Lübeck, German Center for Lung Research (DZL), Lübeck, Germany

## Abstract

The onset of severe SARS-CoV-2 infection is characterized by the presence of afucosylated IgG1 responses against the viral spike (S) protein, which can trigger exacerbated inflammatory responses. Here, we studied IgG glycosylation after BNT162b2 SARS-CoV-2 mRNA vaccination to explore whether vaccine-induced S protein expression on host cells also generates afucosylated IgG1 responses. SARS-CoV-2 naive individuals initially showed a transient afucosylated anti-S IgG1 response after the first dose, albeit to a lower extent than severely ill COVID-19 patients. In contrast, previously infected, antigen-experienced individuals had low afucosylation levels, which slightly increased after immunization. Afucosylation levels after the first dose correlated with low fucosyltransferase 8 (FUT8) expression levels in a defined plasma cell subset. Remarkably, IgG afucosylation levels after primary vaccination correlated significantly with IgG levels after the second dose. Further studies are needed to assess efficacy, inflammatory potential, and protective capacity of afucosylated IgG responses.

**One sentence summary:** A transient afucosylated IgG response to the BNT162b2 mRNA vaccine was observed in naive but not in antigen-experienced individuals, which predicted antibody titers upon the second dose.

## Introduction

Immunoglobulin G (IgG) antibodies (Abs) are crucial for protective immunity in coronavirus disease 2019 (COVID-19) through both fragment antigen binding (Fab)-mediated neutralization and fragment crystallizable (Fc)-mediated effector functions. The IgG Fc-mediated effector functions mainly depend on IgG subclass and Fc *N*-glycosylation, of which the latter has been shown to be important for COVID-19 disease exacerbation (*1–4*). Human IgG contains a single, conserved biantennary *N*-linked glycan at N297 of the Fc portion. This *N-*glycan has a common pentasaccharide core that can further be modified with a fucose, a bisecting *N*-acetylglucosamine (GlcNAc), as well as one or two galactose residues, of which each can further be capped by a sialic acid. Of these glycan residues, galactose and fucose have been described to modulate the activity of complement or natural killer (NK) and myeloid cell IgG-Fc gamma receptors (FcγR), respectively (*5–7*) **(Fig. 1A).**

**Figure 1.**
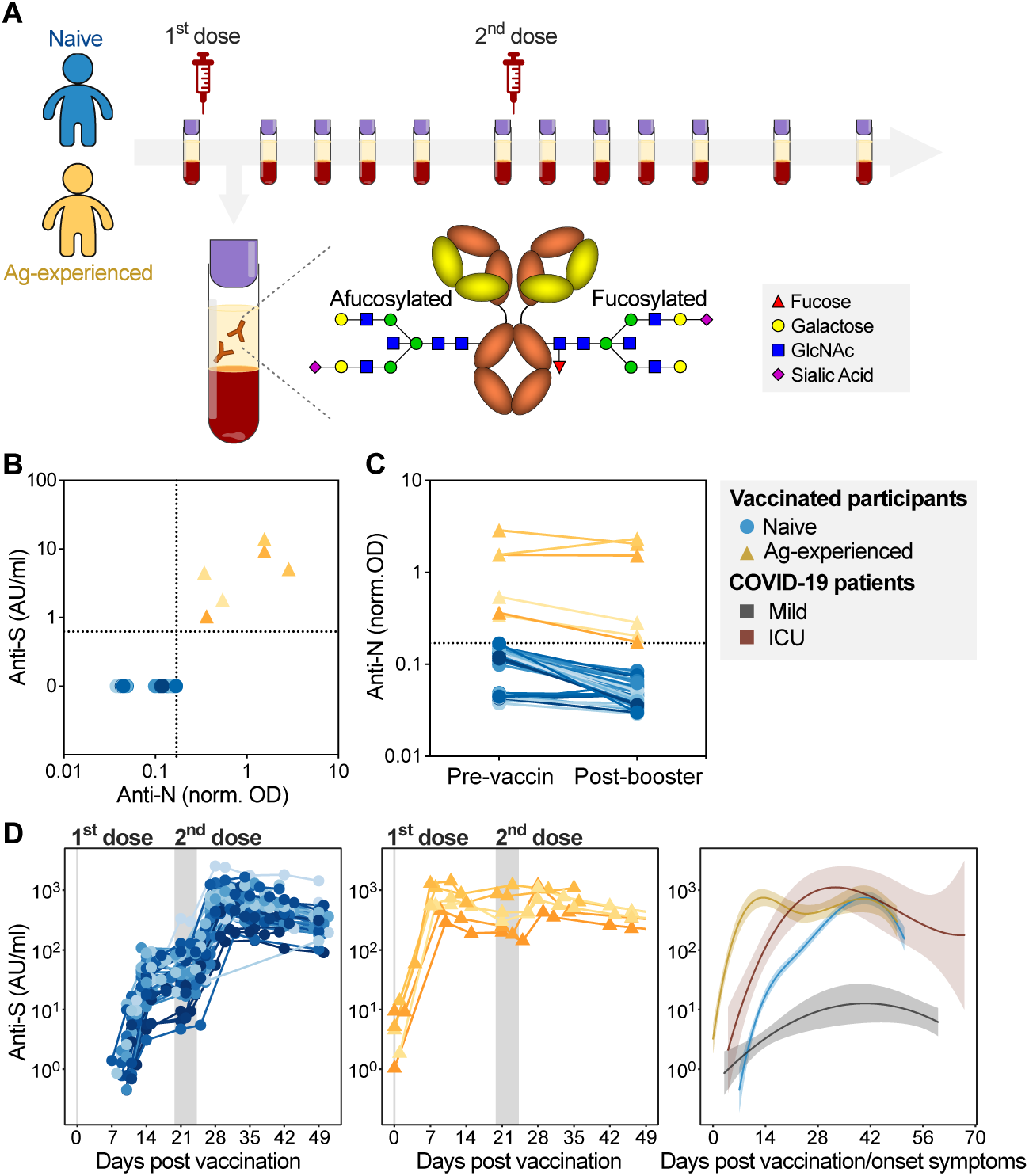
Naive and antigen-experienced individuals show divergent responses to the BNT162b2 mRNA vaccine. **(A)** Schematic depiction of extensive sampling of SARS-CoV-2 naive (blue) and antigen-experienced (yellow) prior to and after the first and second dose of the BNT162b2 mRNA vaccine. **(B-D)** SARS-CoV-2 naive (blue circles) and antigen-experienced vaccines (yellow triangle) IgG levels against **(B)** anti-spike (S) and anti-nucleocapsid (N) IgG levels prior to vaccination and **(C)** anti-nucleocapsid IgG levels during the sampling period. **(D)** Longitudinal anti-S IgG levels for naive (left, cohort 1 (n=33) and 2 (n=9)) and antigen-experienced (middle, cohort 1 (n=6) and 2 (n=0)) vaccinees and corresponding dynamics in comparison to mild (grey) and ICU hospitalized (red) COVID-19 patients (right). Similar data for cohort 3 and 4 are plotted in Fig. S1 and S5, respectively.

Fc-galactosylation levels are highly variable (40-60%), with decreased levels being found in inflammatory diseases such as various infectious, cardiovascular, and autoimmune diseases as well as cancer (*8–12*), whereas increased Fc-galactosylation has been shown to characterize IgG after vaccination (*13, 14*) and COVID-19 infection (*2–4*). Elevated Fc-galactosylation promotes IgG Fc-Fc interaction, leading to hexamerization, which enables docking of complement component 1q (C1q), the first component of the classical complement cascade, and ensuing complement activation (*15, 16*).

Afucosylated IgG has an enhanced binding to FcγRIII, resulting in increased cytokine production and cellular responses, such as Ab-dependent cellular phagocytosis (ADCP) and cytotoxicity (ADCC) (*6, 7, 17*). In healthy conditions, the majority of IgG found in plasma is fucosylated (∼94%) (*18, 19*), but afucosylated, antigen-specific IgG responses have been described in various pathologies, including alloimmune responses to blood cells (*20–22*), as well as immune responses to *Plasmodium (P) falciparum* antigens expressed on erythrocytes (*23*) and to foreign proteins of enveloped viruses, including human immunodeficiency virus (HIV) (*24*), dengue virus (*25*), and severe acute respiratory syndrome coronavirus 2 (SARS-CoV-2) (*2, 3*). The common characteristic of such responses is that the corresponding pathogen-specific antigens are generally expressed on the host cell membrane, unlike most foreign antigens. Intriguingly, pathogen-specific afucosylated IgG1 responses seem to be protective in malaria (*23*) and HIV (*24*), but can, in turn, cause massive inflammation via FcγRIII-mediated pathologies in patients with severe dengue fever (*25*) and have been shown to precede severe COVID-19 (*1–3, 26*).

Non-enveloped viruses, bacteria, and soluble protein-subunit vaccines, which all lack the host cell membrane context, induce almost no afucosylated IgG responses. This includes those of recombinant hepatitis B virus (HBV) and *P. falciparum*-proteins. On the contrary, when expressed in their natural context on host cells, afucosylated IgG responses have been observed in HBV and malaria (*2, 23*). This led us to the hypothesis that antigen presentation at the surface of host cells, possibly together with host co-factors, is required for the induction of afucosylated IgG responses (*2*).

The new mRNA- and adenovirus-based SARS-CoV-2 vaccines induce host cell production of the SARS-CoV-2 spike (S) protein and its subsequent presentation on the cell membrane, unlike traditional soluble protein-subunit vaccines (*27*). Similar to attenuated enveloped-viral vaccines (*2*), mRNA- and adenoviral-based vaccines might therefore also induce an afucosylated IgG response.

Here, we investigated anti-S IgG glycosylation in both naive and antigen-experienced participants after the first and second dose of the BNT162b2 mRNA vaccine against SARS-CoV-2. Additionally, we evaluated glycosyltransferase expression in antigen-specific IgG^+^ plasma cell (PC) subsets to obtain insights into the generation of anti-S IgG glycosylation phenotypes. We furthermore studied the potential contribution of anti-S IgG afucosylation to inflammatory responses using an *in vitro* macrophage activation assay.

## Results

### BNT162b2 mRNA vaccination induces transient afucosylated anti-S IgG in naive, but not antigen-experienced individuals

To analyze the immune response in naive and antigen-experienced individuals upon vaccination with the mRNA vaccine BNT162b2, blood samples were collected from healthy donors at four locations: 1) the Amsterdam University Medical Center (UMC) in The Netherlands, 2) the Fatebenefratelli-Sacco University Hospital in Milan in Italy, 3) the University Medical Center of Schleswig-Holstein Lübeck in Germany, and 4) the Dutch blood bank Sanquin in The Netherlands (**Fig. 1A and Table S2-5**).

To identify antigen-experienced individuals, anti-nucleocapsid (N) and anti-spike (S) IgG responses were investigated both prior to the first dose and during the study, together with previous positive SARS-CoV-2 PCR results (**Fig. 1B-C, S1 and Table S2-5**). Vaccinated SARS-CoV-2 naive individuals showed a detectable anti-S IgG response around day ten after vaccination that further increased upon the second dose (**Fig. 1D and S1A, D, F)**. All vaccinated antigen-experienced individuals had anti-S IgG Abs before vaccination and levels increased fast upon the first dose of BNT162b2 (**Fig. 1D and Fig. S1A, D**). Both naive and antigen-experienced reached similar anti-S levels, which were dominated by IgG1 and IgG3 subclasses against both the S1 and S2 subunits of the S protein (**Fig. S1G**) (*28, 29*). For clarification, we have compared the vaccine-induced responses with the dynamics of mild and intensive care unit (ICU)-admitted COVID-19 patients as described by Larsen *et al*. (*2*) **(Fig. 1D)**.

Next, we explored anti-S and total IgG1 Fc *N*-glycosylation patterns over time (**Fig. 2 and S2-3)**. In both naive and antigen-experienced individuals, an initial drop of anti-S IgG1 bisection levels were seen, with lowered levels as compared to total IgG1 (**Fig. 2A and S2A, 3A**). An early response of highly galactosylated and sialylated anti-S IgG1 was observed in both naive and antigen-experienced individuals, both after the first and second dose (**Fig. 2B-C and S2B-C, S3B-C).** The anti-S IgG1 galactosylation level and time course were similar to what we previously observed in naturally infected individuals with mild symptoms. In contrast, anti-S IgG1 galactosylation has been shown to drop rapidly in ICU-admitted COVID-19 patients, as previously described **(Fig. 2B)** (*2, 30*). A high level of IgG galactosylation boosted the classical complement pathway activation capacity through enhanced C1q-binding, which was in line with previous reports **(Fig. S4)** (*15*). Anti-S IgG1 sialylation follows the galactosylation trend, with an increase after the first and second dose **(Fig 2C, S2C)**.

**Figure 2.**
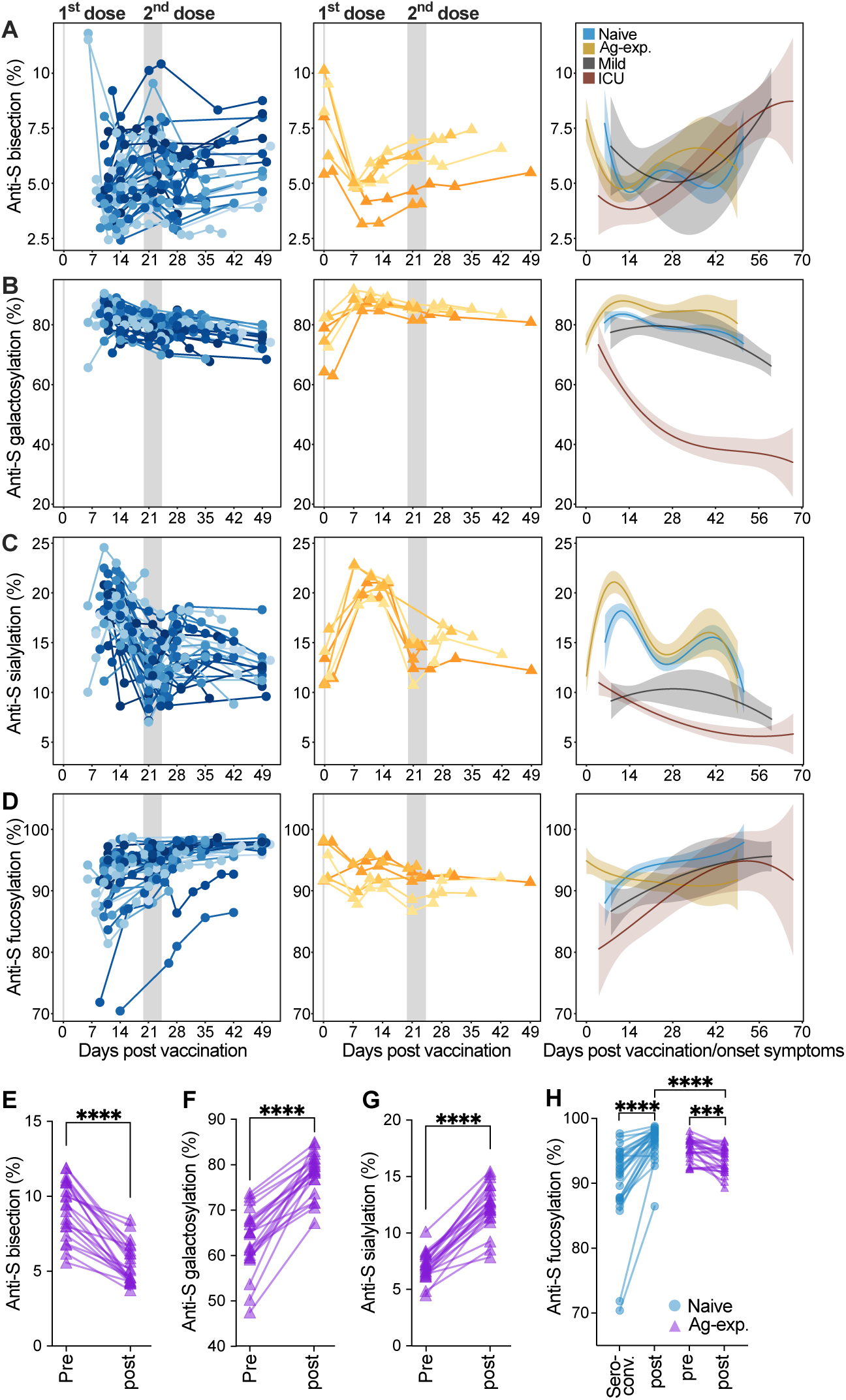
Anti-Spike IgG1 glycosylation is dynamic. Longitudinal anti-S IgG1 Fc **(A)** bisection, **(B)** galactosylation, **(C)** sialylation, and **(D)** fucosylation for naive (left, blue, cohort 1 (n=33) and 2 (n=9)), antigen-experienced (middle, yellow, cohort 1 (n=6) and 2 (n=0)) in comparison to mild (grey) and ICU hospitalized (red) COVID-19 patients (right) anti-S IgG1 galactosylation from our previous study (*2*). Anti-S IgG1 Fc **(E)** bisection, **(F)** galactosylation, **(G)** sialylation, and **(H)** fucosylation for the additional vaccinated antigen-experienced plasma donors (purple, cohort 4 (n=22)) before (pre) and after (post) vaccination and **(H)** in comparison to naive vaccinees at seroconversion (seroconv.) and after the second dose (post) (blue, cohort 1 (n=33) and 2 (n=9)). Differences were assessed using the Mann-Whitney U test (***, ****: *p*-value < 0.001, 0.0001, respectively). Similar data for cohort 3 are plotted in Fig. S2.

We recently hypothesized that afucosylated IgG, hardly seen in responses to soluble protein or polysaccharide antigens, are specifically induced against foreign antigens on host cells (*2*). In agreement with this, up to 25% of anti-S IgG1 Fc was found to be afucosylated after vaccination with the BNT162b2 SARS-CoV-2 mRNA vaccine, in comparison to *∼*6% of afucosylated total IgG1 found in serum or plasma (**Fig. 2D and S2D, S3D**). This pronounced afucosylation pattern was observed only early on in naive individuals after the first dose of BNT162b2, which gradually decreased to levels similar to total IgG1 at four weeks post seroconversion (**Fig. 2D and S2D, S3D**). This early, transient afucosylated response in naive vaccinees after the first dose was less prominent when compared to ICU-admitted COVID-19 patients and most individuals with mild symptoms (**Fig. 2D**).

In contrast, antigen-experienced individuals had an anti-S IgG1 afucosylation level of ∼2-10% and slightly increased after vaccination (**Fig. 2D, S2D**). We further investigated this by expanding the vaccinated antigen-experienced through recruiting vaccinated blood donors previously infected with SARS-CoV-2 **(Fig. S5, Table S5)**. Similar anti-S IgG1 Fc bisection, galactosylation, sialylation, and fucosylation dynamics were observed (**Fig. 2E-H)**. Compared to naive individuals, this antigen-experienced cohort showed a significantly lower anti-S IgG1 Fc fucosylation after vaccination (**Fig. 2H**). No temporal changes were observed for total IgG glycosylation (**Fig. S2-3).**

### Impact of afucosylated anti-S IgG is limited because of low antibody levels

We assessed the effector function of the anti-S Abs induced by BNT162b2 mRNA vaccination by testing their capacity to induce macrophage-driven inflammatory responses. For this, we measured IL-6 production by human-derived, *in vitro* differentiated, alveolar-like macrophages. These were stimulated overnight by exposure to immune complexes (ICs) generated from S protein and vaccinees’ sera in the presence and absence of virus-like co-stimuli (polyinosinic:polycytidylic acid (poly(I:C))) (**Fig. 3A**) (*31*). Notably, anti-S ICs from antigen-experienced individuals induced significantly higher IL-6 levels compared to naive individuals for all time points after the first dose, with the most pronounced difference seen at day 10 (**Fig. 3B**). IL-6 induction was similar for both groups after the second dose as IgG levels became comparable (**Fig. 1D, 3B and S6**). Despite the clear difference between both groups, IL-6 levels were relatively low for all conditions, which is in line with various previous findings showing that IgG ICs only induce pro-inflammatory cytokine production in the presence of both high afucosylation and antibody levels in the presence of viral or bacterial co-stimulus that activates receptors such as Toll-like Receptors (TLRs) (*1, 32, 33*).

**Figure 3.**
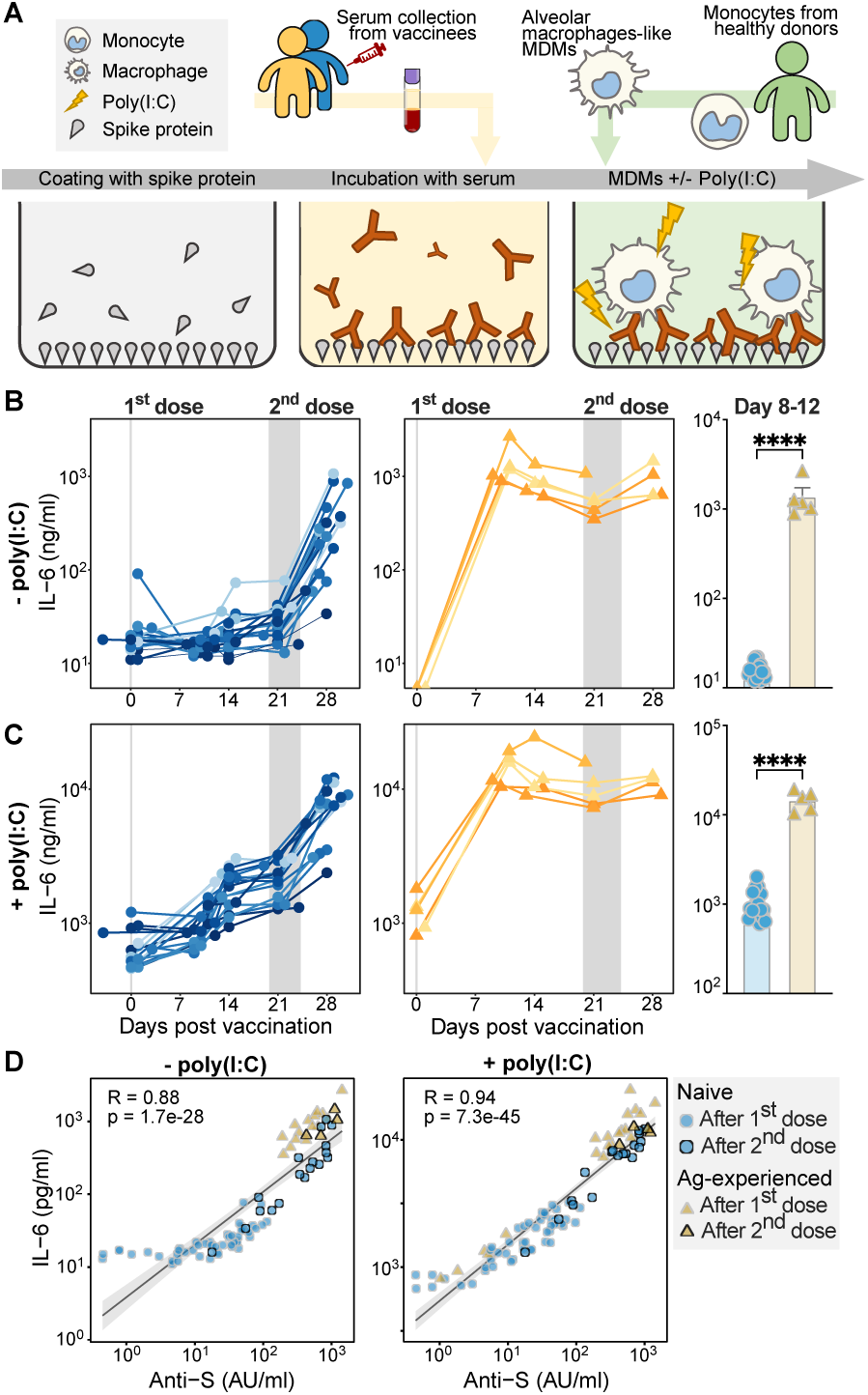
Antibody levels are primarily responsible for macrophage activation. **(A)** Schematic representation of the alveolar-like monocyte-derived macrophages stimulation assay with and without polyinosinic:polycytidylic acid (poly(I:C)). **(B-C)** IL-6 responses of macrophages stimulated with spike protein and naive (left, blue) and antigen-experienced (middle, yellow) vaccinee sera and a comparison of IL-6 response between day 8 to 12 by unpaired *t*-test (right) in the **(B)** absence or **(C)** presence of poly(I:C). **(D)** Correlation between IL-6 levels and anti-S IgG levels in the absence (left) and presence (right) of poly(I:C) stimulation. All data represent a subgroup of cohort 1 (n=23, see Table S2).

To further test the inflammatory capacity of anti-S IgG, we also measured IL-6 production upon TLR co-stimulation with the TLR3 ligand poly(I:C). Upon TLR co-stimulation, anti-S ICs strongly amplified IL-6 production by human macrophages in both groups **(Fig. 3C)**. Again, the difference between the vaccinated naive and antigen-experienced individuals was most pronounced around day ten post vaccination (**Fig. 3C**). In both cases, the capacity of the sera to activate these macrophages seem to be explained by antibody levels (**Fig. 3D**). However, when anti-IgG levels became comparable after the second dose, the sera of antigen-experienced individuals induced only slightly higher IL-6 levels both with and without poly(I:C) (**Fig. 3B-D and S6**), which correlated with higher afucosylation levels, but not with other IgG1 glycosylation traits (**Fig. S7**). Combined, these data suggest that the transient afucosylated anti-S IgG that is produced after vaccination of naive individuals has little effect on macrophage activation, because it is accompanied by low antibody levels.

### Differential plasma cell responses in naive and antigen-experienced individuals

In line with literature, a highly sialylated IgG glycosylation phenotype was observed early after vaccination, regardless of antigen experience (**Fig 2C and S2C, 3C**) (*14, 34, 35*). Interestingly, this early, transient, high sialylation was particularly pronounced for fucosylated anti-S IgG1, for both naive and antigen-experienced vaccinees until day fourteen after the first dose (**Fig. 4A and S8**). Anti-S IgG1 Fc galactosylation levels of neither naive nor antigen-experienced showed a difference between fucosylated and afucosylated anti-S IgG1 (**Fig. 4B and S8**). This result led to the hypothesis that early highly galactosylated and sialylated anti-S IgG1 and afucosylated anti-S IgG1 might be produced by different PC subsets.

**Figure 4.**
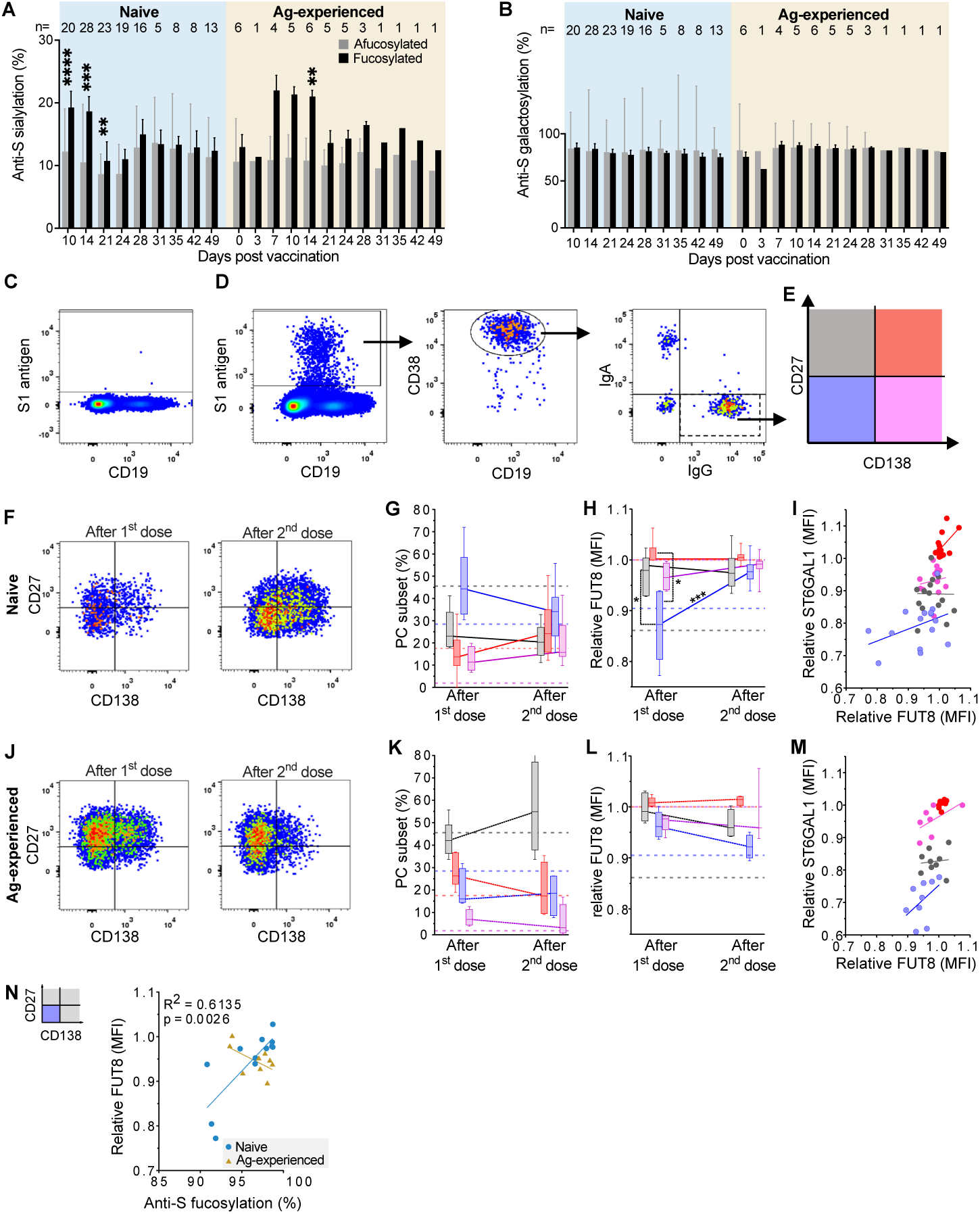
Different populations of B cells express distinct levels of glycosyltransferases. **(A)** Sialylation and **(B)** galactosylation levels of afucosylated (grey) and fucosylated (black) anti-S IgG for naive (left) and antigen-experienced (right) vaccinated participants over time of cohort 1 (n=39) and 2 (n=9) after re-normalization by setting the sum of all afucosylated glycoforms to 100% and all fucosylated glycoforms to 100%. Glycosylation levels were compared by a paired t-test. **(C-N)** Flow cytometry analysis of blood cells gated on single, living lymphocytes from naive and antigen-experienced vaccinees (subset of cohort 3 (n=15), see Table S2) were analyzed 7-14 days upon the first (naive: n=6 and antigen-experienced: n=5) or 5-8 days upon the second (naive: n=15 and antigen-experienced: n=4) dose. **(C-E)** Gating strategy exemplified for a naive individual **(C)** pre-immunization and **(D)** after the first dose. S1-reactive B cells were gated and further gated for CD19^int^ CD38^+^ PCs to analyze IgG^+^ PC subsets as defined by **(E)** CD27 and CD138. **(F-G)** Naive and **(J-K)** antigen-experienced vaccinees analyzed according to the gating strategy. **(H, L)** Relative fucosyltransferase 8 (FUT8; median (MFI)) expression per IgG^+^ PC subset and **(I, M)** its correlation with relative alpha2,6-sialyltransferase (ST6GAL1) expression. **(N)** Relative FUT8 expression of CD27^low^CD138^-^ IgG^+^ PCs correlated with anti-S IgG1 Fc fucosylation found in the corresponding serum (**Fig. S9C**). The median (MFI) of FUT8 or ST6GAL1 expression in CD138^+^ IgG+ S1-reactive PCs of each sample was set to 1 for inter-assay comparison. Dotted horizontal lines indicate corresponding values of total IgG^+^ PC subsets from untreated healthy controls (**Fig. S11**).

Next, we analyzed the anti-S1 blood-derived IgG^+^ CD38^+^ PC subset responses to assess whether they phenotypically diverge in their anti-S IgG1 glycosylation pattern between naive and antigen-experienced individuals (*36–38*). We found CD27^low^ CD138^-^ IgG^+^ CD38^+^ PCs to be dominant in naive individuals after both the first and second dose (**Fig. 4C-G, S9A-G, S10A-F**). In contrast, antigen-experienced vaccinees primarily induced CD27^+^ CD138^-^ IgG^+^ CD38^+^ PCs after both doses (**Fig. 4J-K, S9A-I, S10A-F)**, which was also the dominant subset in total IgG^+^ PCs of the naive antigen unvaccinated controls **(Fig. S10A-F, S11**).

We found that *α*1,6-fucosyltransferase 8 (FUT8; the glycosyltransferase responsible for core fucosylation (*39*)) protein expression was lowest in the CD27^low^ CD138^-^ IgG^+^ PC subset in naive individuals after the first, but not the second dose (**Fig. 4H-I, L-M and S10H**). The α2,6-sialyltransferase 1 (ST6GAL1; the glycosyltransferase responsible for α2,6-linked sialylation (*35*)) protein expression was highest in the CD27^+^ CD138^+^ and the lowest in CD27^low^ CD138^-^ IgG^+^ PC subset after both doses in naive and antigen-experienced individuals, as well as in total IgG^+^ PCs of unvaccinated healthy control individuals (**Fig. 4I, M and S10G, S11**). In naive individuals, FUT8 expression in CD27^low^ CD138^-^ IgG^+^ PCs correlated with anti-S IgG1 fucosylation (**Fig. 4N**).

### Early afucosylated anti-S correlates with anti-S IgG titer upon the second dose

Thorough examination of individual study participants revealed that vaccinees with high afucosylated anti-S IgG1 often show high anti-S IgG titers **(Fig. S12)**. To study this possible link, we correlated the anti-S IgG1 Fc glycosylation both upon seroconversion **(Fig. 5A-D and S13B-E)** and after the first dose **(Fig. 5E-H and S13F-I)** with the anti-S IgG levels after the first **(Fig. S13B-I)** and second dose **(Fig. 5)**. For this, we selected the anti-S IgG level on the day of the second dose up to three days prior, and the highest level reached up to two weeks post the second dose, respectively. Even though anti-S IgG levels after the first dose correlated with the levels after the second dose **(Fig. S13A)**, anti-S IgG1 Fc afucosylation either at seroconversion or at the time of the second dose correlated with the IgG levels after the second dose **(Fig.5D, H)** but not with the first dose **(Fig S13E, I)**. No correlations were found for IgG levels and the other anti-S IgG glycosylation traits **(Fig 5A-C, E-G, and S13B-D, F-H)**.

**Figure 5.**
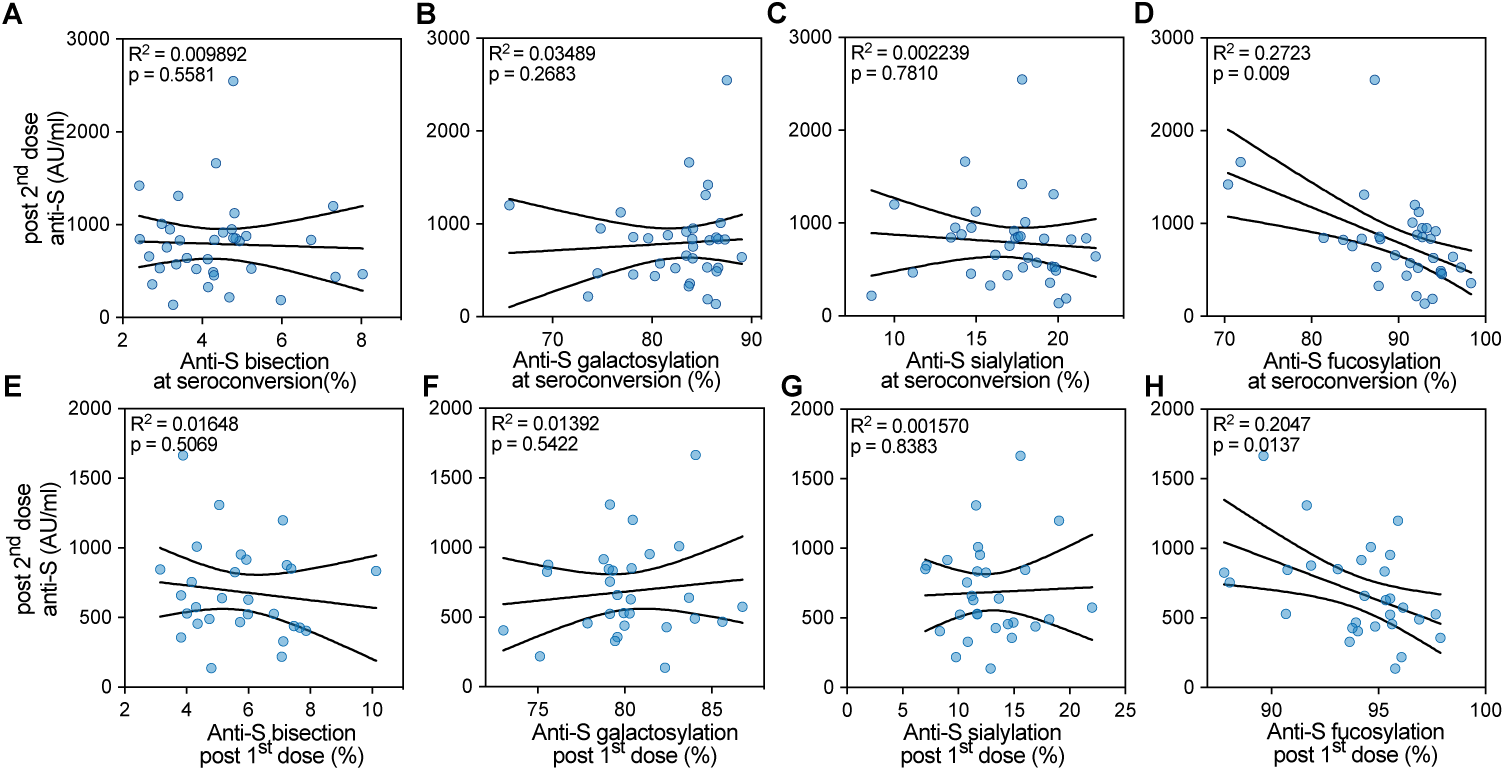
Afucosylation of anti-S IgG1 correlates with titer after the second dose. Correlation analysis of anti-S IgG1 Fc glycosylation with levels for naive vaccines from cohort 1 (n=33) and 2 (n=9) **(A-D)** Correlation of anti-S IgG1 Fc glycosylation upon seroconversion and **(E-H)** later after the first dose (on the day of the second dose up until three days prior) with levels after the second dose (highest levels up to 14 days after 2^nd^ dose).

## Discussion

The mRNA vaccine-induced presentation of the viral S protein on the membrane of host cells mimics the S protein presentation during natural SARS-CoV-2 infections. Enveloped viruses, attenuated enveloped viral vaccines, but also *P. falciparum*-infected erythrocytes and alloantigens on blood cells express antigens on host cells and induce persistent afucosylated IgG responses (*2, 22–25, 40*). Although we previously found afucosylated IgG responses to be associated with strong pro-inflammatory responses in critically ill COVID-19 patients (*1, 2*), this type of response seems to be protective in HIV infections (*24*) and malaria (*23*).

Here, we show for the first time that the BNT162b2 mRNA vaccine induces afucosylated anti-S IgG1 responses in SARS-CoV-2 naive individuals upon seroconversion, which decreases within four weeks to the level of total IgG1. Recent work from Farkash *et al.* and Chakraborty *et al.* did not pick up this transient response due to sampling of two and four weeks after the first dose, respectively (*26, 41*). This afucosylated response was similar, but less pronounced than observed in natural SARS-CoV-2 infections (*2, 30*). The transient afucosylated IgG1 glycosylation pattern after vaccination suggests that a co-stimulus may be missing to induce memory B cells and long-lived IgG^+^ PCs producing stable anti-S IgG1 afucosylation levels. Alternatively, a missing local type of inflammatory signal might provide a negative feedback steering developing B cells to produce fucosylated IgG. In contrast, SARS-CoV-2 antigen-experienced vaccinees start off with low (∼2-10%), but persistent anti-S IgG1 afucosylation levels which slightly increase upon vaccination, assuming re-activation of memory B cells generating afucosylated IgG antibodies.

Our analyses revealed that the differences in the effector functions elicited by anti-S ICs on macrophages between naive and antigen-experienced vaccinees mainly depend on the titer after the first dose, with afucosylation only being a secondary factor. Our previous work has shown that exaggerated pro-inflammatory responses were only observed with serum containing high titers of considerably afucosylated IgG1 (>10%) (*1, 2*). Such afucosylated IgG1 levels in this study were only observed early after seroconversion in naive individuals and not in combination with high titers. In line with this, anti-S ICs from vaccinee’s sera induced very moderate pro-inflammatory cytokine levels in the absence of TLR co-stimulation, suggesting low inflammatory side effects in both groups after immunization with the BNT162b2 mRNA vaccine. Nevertheless, when comparing naive and antigen-experienced vaccinees after the second dose, when anti-S IgG levels were comparable, antigen-experienced individuals induced slightly higher IL-6 production in the macrophage activation assay, which is in agreement with their higher levels of afucosylated IgG.

Moreover, the different immune responses of naive versus antigen-experienced individuals upon vaccination were reflected in the antigen-specific PC response. Whereas naive individuals primarily induced CD27^low^ CD138^-^ IgG^+^ CD38^+^ PCs, antigen-experienced individuals primarily induced CD27^+^ CD138^-^ IgG^+^ CD38^+^ PCs after both doses. Furthermore, only the naive subpopulation showed reduced FUT8 expression in CD27^low^ CD138^-^ IgG^+^ CD38^+^ PCs only after the first vaccination, which correlated with the amount of afucosylated IgG1 observed in these individuals (*26*). The existence of an IgG^+^ PC subset responsible for the biosynthesis of afucosylated IgG Abs in antigen-experienced individuals has yet to be identified in further studies.

In accordance with previous reports on immunization (*14, 34, 35*) and Fc glycosylation after BNT162b2 mRNA vaccination (*26, 41*), we also observed transiently, highly sialylated anti-S IgG1 at one to two weeks after both the first and second dose, which has been suggested to facilitate antigen presentation in subsequent GC reactions for improving affinity maturation (*14, 42*). Furthermore, the anti-S IgG1 for both naive and antigen-experienced vaccinees was extensively galactosylated. These high levels of IgG galactosylation have been shown to boost the capacity to activate the classical complement pathway, through enhanced C1q-binding, in line with recent findings that have shown galactosylation promotes IgG1 hexamerization ultimately leading to increased C1q-binding and ensuing classical complement activation (*15, 16, 41*).

Afucosylated IgG1 may support antigen presentation on antigen-presenting cells through FcγRIIIa, as well as inducing better T-helper and memory B cell responses, and subsequent booster responses (*43–45*). In support of this, we observed that early afucosylated anti-S IgG1 responses correlated significantly with anti-S IgG levels after the second dose in naive individuals (*41*). At seroconversion, afucosylated anti-S IgG1 Abs in naive vaccinees might provide enhanced protection, even without high titers. Over time, when afucosylated IgG levels drop, protection in these individuals might be compensated by the increased anti-S IgG levels, which should be considered for the timing of subsequent vaccination. Furthermore, reduced levels of anti-S IgG1 afucosylation might reduce the risk of pro-inflammatory side effects, with a trade-off of dampened Fc-mediated effector functions upon pathogen contact. In antigen-experienced individuals, matters are reversed, as these individuals start off with lower afucosylated anti-S IgG levels prior to vaccination, which significantly increased after vaccination. This suggests an enhanced corresponding memory B cell response, which would be in line with stronger protection in this group (*46–48*). Similarly, a gradual increase in afucosylation has been observed with repeated natural immunizations to antigens displayed on the membrane of *P. falciparum*-infected red blood cells (*23*). This is in contrast to alloimmunization to the red blood cell RhD antigen, where the afucosylated response in hyperimmune donors is very stable over time (*40*). The increased level of afucosylated anti-SARS-CoV2 IgG in vaccinated antigen-experienced individuals might have a positive impact on the therapeutic effect of convalescent plasma, as especially these donors are presently selected for clinical trials and it has been shown that increased ADCC activity of the administered antibodies is positively correlated with outcome.

Due to the limited sample size, we did not stratify study participants according to sex and age, which may influence IgG glycosylation profiles (*18*). However, outside of the context of a specific pathology, total IgG fucosylation levels remain constant throughout life with the exception of an initial decrease after birth (*19*). A second limitation of our study is the uneven sample size for naive and antigen-experienced vaccine recipients after the first and second dose of BNT162b2. This is largely due to lack of an accessible, high-throughput serological assay to measure antigen-specific IgG glycosylation to study both transient and stable glycosylation features in disease settings.

In summary, our data demonstrate a qualitatively and quantitatively distinct IgG immune response between BNT162b2 mRNA vaccinated SARS-CoV-2 naive and antigen-experienced individuals. Transient afucosylated IgG1 responses were induced in naive individuals upon the first dose, which correlated with increased titer after the second vaccination. In contrast, antigen-experienced vaccinees had low levels of afucosylated anti-S, which slightly increased upon vaccination. The qualitatively distinct IgG1 glycosylation patterns might further mediate differences in protection between these two groups. Future efforts focused on inducing and studying antigen-specific, afucosylated IgG1 responses are needed to investigate their protective capacity and inflammatory potential in anti-viral and vaccine-induced immunity.

## Materials and Methods

This study was designed to investigate the effect of the BNT162b2 BioNTech/Pfizer mRNA vaccine on anti-Spike IgG1 Fc glycosylation and PC subsets. We obtained serum, plasma and/or PBMC samples from vaccinated participants from 1) healthcare works at the Amsterdam UMC, The Netherlands (n=39), 2) The Fatebenefratelli-Sacco Infectious Diseases Physicians Group (n=9), 3) the University Medical Center of Schleswig-Holstein, Lübeck, Germany (n=40), and 4) the Dutch blood bank Sanquin, The Netherlands. The discrimination between vaccinated SARS-CoV-2 naive and antigen-experienced participants was made by serology (anti-Spike and anti-Nucleocapsid IgG) and positive PCR-tests before vaccination. No other selection criteria were used and participants were selected at random.

### Vaccination study cohorts and control individuals

#### Cohort 1. Amsterdam UMC cohort

Subjects were part of the S3 cohort study (S3 cohort; NL 73478.029.20, Netherlands Trial Register NL8645), a prospective serologic surveillance cohort study among hospital healthcare workers in the Amsterdam University Medical Center (Amsterdam UMC). Between January and March 2021, 39 cohort participants received their first dose of BioNTech/Pfizer mRNA vaccine (BNT162b2, 30ug) (**Table S2**). A second dose was administered approximately 21 days after the first dose. Samples were obtained directly before and 3, 7, 10 and 14 days after the first dose, and directly before and 3, 7, 10, 14, 21 and 28 days after the second dose (**Table S2**). Participants were included through informed consent. The ethics committee of the AUMC approved the study.

#### Cohort 2. The Fatebenefratelli-Sacco Infectious Diseases Physicians Group

Nine healthcare workers at the Luigi Sacco Infectious Diseases Hospital, Milano, Italy were immunized with BioNTech/Pfizer mRNA vaccine (BNT162b2, 30ug) and received a 2^nd^ dose 21 days after the 1^st^ dose. Blood samples were obtained directly before the 1^st^ dose, and twice a week for six weeks from December 2020 to February 2021 after obtaining informed consent (**Table S3**).

#### Cohort 3. Lübeck cohort

Forty subjects were recruited at the University Medical Center of Schleswig-Holstein, Lübeck, Germany from December 2020 (including samples (participants 1-22; **Table S4**) described in Lixenfeld et al. (*28*)): 1) 32 individuals immunized with the BioNTech/Pfizer vaccine BNT162b2 (30 µg) without or with known SARS-CoV-2 infection history (19 of these 32 individuals (analyzed in Fig. S1) received the 2^nd^ dose between day 32 and 37 after the 1st) and 2) and 8 unvaccinated individuals without SARS-CoV-2 infection history as negative control (**Table S4**). Blood samples were collected after obtaining written informed consent under the local ethics board–approved protocols 19-019(A) and 20-123 (Ethics Committee of the University of Lübeck, Germany).

#### Cohort 4. Convalescent plasma donors

Sanquin blood donors (n=22) found seropositive for SARS-CoV-2 prior to vaccination were included in the study (**Table S5**). All participants provided written informed consent. The study was approved by the Academic Medical Center Institutional Medical Ethics Committee of the University of Amsterdam.

All studies complied with the latest version of the Declaration of Helsinki.

### Anti-SARS-Cov2 Ab levels

#### Cohort 1, 2 and 4

Anti-S IgG Abs levels were measured by coating MaxiSorp NUNC 96-well flat-bottom plates (Thermo Fisher Scientific, Roskilde, Denmark) overnight with 1 µg/ml recombinant, in-house produced trimerized spike protein in PBS, as described before (*49*). The following day, plates were washed five times with PBS supplemented with 0.02% polysorbate-20 (PBS-T) and incubated for 1 hour with a dilution range of plasma from the Amsterdam UMC cohort in PBS-T supplemented with 0.3% gelatin (PTG). A serially diluted plasma pool, obtained by combining plasma from a collection of convalescent COVID-19 donors (*50*), was used as a calibrant. After incubation, plates were washed five times with PBS-T and incubated with 1 µg/ml anti-human IgG-horseradish-peroxidase (HRP) (clone: MH16.1, Sanquin, Amsterdam, the Netherlands). After washing, Ab binding was evaluated by adding 50% diluted tetramethylbenzidine substrate (1-step ultra TMB, #34029, Thermo Scientific). The reaction was terminated by adding equal amounts of 0.2 M H_2_SO_4_ (Merck, Darmstadt, Germany) and absorbance was measured at 450 and 540 nm. The calibrant plasma pool was assigned the value of 100 arbitrary units (AU), which corresponds to approximately 21 µg/ml (*51*).

Anti-Receptor Binding Domain (RBD) and anti-Nucleocapsid (N) antibody levels were measured as by an RBD and N-based bridging assay, respectively, as described previously (*50, 51*).

#### Cohort 3

To detect anti-S1 IgG as well as anti-NCP IgG Abs, serum samples were collected on the indicated days (**Table S4**) and EUROIMMUN SARS-CoV-2 S1 IgG (EUROIMMUN, Luebeck, Germany; #EI 2606-9601-2 G) and EUROIMMUN SARS-CoV-2-NCP IgG (#EI 2606-9601-2 G) ELISA were performed according to the manufacturer’s instructions, respectively.

To detect anti-S1 and -S2 IgG and IgG subclass (IgG1-4) Abs, 96-well ELISA plates were coated alternatively with 4 µg/ml of SARS-CoV-2-S1 (ACROBiosystems, Newark, DE 19711, USA; #S1N-C52H3) or -S2 (ACROBiosystems; #S2N-C52H5) antigen per well (HL-1 ELISA (*28*)). The plates were washed with PBS-T. Subsequently, sera (diluted 1/100 or 1/1000 in 0.05% Tween-20, 3% BSA in PBS) were added. Bound Abs were detected with HRP-coupled polyclonal goat anti-human IgG Fc (#A80-104P)-specific Abs purchased from Bethyl Laboratories (Montgomery, TX, USA), or monoclonal anti-human IgG1 (clone HP-6001), IgG2 (clone HP-6014), IgG3 (clone HP-6050), or IgG4 (clone HP-6025)-specific Abs purchased from Southern Biotech (Birmingham, AL, USA) in 0.05% Tween 20, 3% BSA in PBS. After incubation with the tetramethylbenzidin (TMB) substrate (BD Biosciences, San Diego, CA, USA) and terminating of the reaction with the addition of H_2_SO_4_, the optical density (OD) was measured at 450 nm. The specificities of the secondary Abs have been verified recently (*28*).

### IgG Fc glycosylation analysis by mass spectrometry

Anti-S IgG Abs were affinity-captured from plasma or sera using recombinant, in-house produced trimerized spike protein-coated plates (Thermo Fisher Scientific, Roskilde, Denmark) followed by a 100 mM formic acid elution step, as described elsewhere (*2, 49*). Total IgG Abs were affinity-captured from plasma or sera using a Protein G AssayMAP Cartridge Rack on the Bravo (Agilent Technologies, Santa Clara, CA) or Protein G Sepharose 4 Fast Flow beads (GE Healthcare, Uppsala, Sweden) in a 96-well filter plate (Millipore Multiscreen, Amsterdam, Netherlands), respectively, as described elsewhere(*2, 30, 52*).

Eluates from both anti-S and total IgG affinity-purification were dried by vacuum centrifugation and subjected to tryptic cleavage followed by LC-MS analysis as described previously (*2, 30*).

### LC-MS data processing and method robustness

Raw LC-MS spectra were converted to mzXML files. LaCyTools, an in-house developed software was used for the alignment and targeted extraction of raw data (*53*). Alignment was performed based on average retention time of at least three high abundant glycoforms. The analyte list for targeted extraction of the 2^+^ and 3^+^ charge states was based on manual annotation as well as on literature reports (*2, 54*). Inclusion of an analyte for the final data analysis was based on quality criteria including signal-to-noise (higher than 9), isotopic pattern quality (less than 25% deviation from the theoretical isotopic pattern), and mass error (within a ±20 parts per million range) leading to a final analyte list (**Table S5**). Relative intensity of each glycan species in the final analyte list was calculated by normalizing to the sum of their total areas. Normalized intensities were used to calculate fucosylation, bisection, galactosylation and sialylation (**Table S6**).

### Complement ELISAs

Pierce™ Nickel Coated Clear 96-well plates (Thermo Fisher Scientific, #15442) were incubated with 100 µL of 1 µg/mL purified RBD-protein for 1 hour at RT. Hereafter, the plates were washed five times with 0.05% PBS-Tween20 and incubated with 100 µL glycoengineered COVA1-18 (2C1) hIgG1 mAbs for 1 hour at RT (*1, 49*). A two-fold dilution series was used, with a starting concentration of 20 µg/ml. Subsequently the plates were washed and 100 µL of 1:35 pooled human serum in Veronal Buffer (*5*) with 0.1% poloxamer 407, 2 mM MgCl_2_ and 10 mM CaCl_2_ was added and incubated for 1 hour at RT, as described previously (*15*). Consequently, the plates were washed and 100 µL 1/1000 anti-C1q-HRP (*55–57*) was added and incubated for 1 hour at RT. Lastly, the plates were washed and developed with 100 µL 0.1 mg/mL TMB solution with 0.11M NaAc and 0.003% H_2_O_2._ The reaction was terminated with 100 µL 2M H_2_SO_4_ and the absorbance was measured using the Biotek Synergy™ 2 Multi-Detection Microplate Reader at 450-540 nm.

The binding capacity of the glycoengineered COVA1-18 (2C1) hIgG1 mAbs was tested by directly coating Nunc MaxiSorp flat-bottom 96-well plates (Thermo Fisher Scientific) O/N at 4°C with 100 µL 1 µg/mL purified SARS-CoV-2 RBD-protein. The plates were washed with PBS-T and incubated with 100 µL glycoengineered COVA1-18 (2C1) hIgG1 mAbs for 1 hour at RT. A two-fold dilution series was used, with a starting concentration of 1 µg/ml. Hereafter, the plates were washed and incubated with 100 µl of 1/1000 Mouse Anti-Human IgG Fc-HRP (Southern-Biotech) for 1 hour at RT. Lastly, the plates were washed and developed with TMB solution. The reaction was terminated with 2M H_2_SO_4_ and the absorbance was measured using at 450-540 nm.

### IL-6 ELISA

Supernatants of stimulated alveolar-like monocyte-derived macrophages (MDMs) were harvested after 24 hours to determine cytokine production. IL-6 levels in the supernatant were measured by enzyme-linked immunosorbent assay (ELISA) using IL-6 CT205-c and CT205-d antibody pair (U-CyTech, Utrecht, the Netherlands) as described previously (*1*).

### Alveolar-like monocyte-derived macrophage differentiation

Buffy coats from healthy donors were obtained from Sanquin Blood Supply (Amsterdam, the Netherlands). Monocytes were isolated from buffy coat by density gradient centrifugation using Lymphoprep^TM^ (Axis-Shield, Dundee, Scotland) followed by CD14^+^ selection via magnetic cell separation using MACS CD14 MicroBeads and separation columns (Miltenyi Biotec, Bergisch Gladbach, Germany), as previously described (*31*). Alveolar-like MDMs were generated by differentiating CD4^+^ monocytes on tissue culture plates into macrophages in the presence of 50 ng/ml of human M-CSF (Miltenyi Biotec, Bergisch Gladbach, Germany) for 6 days, followed by 24-hour incubation in culture medium supplemented with 50 ng/ml IL10 (R&D System, Minneapolis, MN, USA). The resulting MDMs were then detached for stimulation using TrypLE Select (Gibco, Waltham, MA).

### Cell stimulation

96-well high affinity plates were coated with 2 µg/ml soluble perfusion stabilized Spike protein as described previously (*1*). After overnight incubation, plates were blocked with 10 % FCS in PBS for 1 hour at 37 °C. Diluted heat-inactivated serum (Table S1, 1:50 dilution) was added for 1 h at 37 °C. 50,000 cells/well were stimulated in the pre-coated plates in culture medium (Iscoves’s Modified Dulbecco’s Culture Medium (IMDM) (Gibco) containing 5% FBS (Capricorn Scientific, Ebsdorfergrund, Germany) and 86 µg/ml gentamicin (Gibco) without or supplemented with 20 µg/ml polyinosinic:polycytidylic acid (poly(I:C)) (Sigma-Aldrich, Darmstadt, Germany).

### Flow cytometric analysis of blood samples

Blood samples were collected at the indicated days in EDTA-tubes and processed or frozen within the next three hours for flow cytometric analysis (Attune Nxt; Thermo Fisher Scientific) of different B cell populations (*28*). Peripheral blood mononuclear cells (PBMCs) were obtained by gradient centrifugation in Ficoll. The following fluorochrome-coupled Abs were used for surface staining: anti-CD19 (Biolegend; clone HIB19), anti-CD38 (Biolegend: HIT2), anti-IgG Fc (Biolegend; M1310G05), anti-CD27 (Biolegend: 0323) and anti-CD138 (Biolegend: MI15) as well as LIVE/DEAD Fixable Near-IR stain (Thermofisher; L34976). For additional intracellular staining, samples were fixed with Cytofix/Cytoperm according to the manufacturer’s instructions (BD Biosciences) followed by permeabilization (0.05% saponin, 0.1% BSA in 0.05 x PBS) and additional staining with anti-IgG, anti-human ST6GAL1 (R&D Systems, polyclonal goat IgG Ab; #AF5924), or isotype goat control IgG (R&D Systems), or anti-human FUT8 (R&D Systems, polyclonal sheep IgG; #AF5768), or isotype sheep control IgG (R&D Systems), as well as SARS-CoV-2-S1 (biotin-coupled; Acro; #S1N-C82E8) and fluorochrome-coupled streptavidin (Biolegend). The anti-ST6GAL1 and anti-FUT8 Abs were labeled with Alexa Fluor 488 labeling kit (Life Technologies GmbH; #A20181). 20 million cells were recorded per sample. Flow Cytometry Standard (FCS) 3.0 files were analyzed with FlowJo software version X 0.7 (BD Biosciences).

### Statistical analysis

The log_10_ values of the anti-spike IgG titers were used for the correlation analyses between log_10_ values of the measured concentrations of IL-6 (in pg/ml). The percentages of anti-S IgG1 glycosylation traits were used for the color overlay. The Pearson correlation coefficient (*R*) and associated *P*-value are stated in each graph. For the comparison of the IL-6 concentration produced by alveolar-like macrophages, an unpaired *t*-test was performed. These analyses were performed in the R statistical environment (v3.6.3).

Other statistical analyses were performed using GraphPad Prism v6.0 (GraphPad, La Jolla, CA). Differences in anti-S IgG1 glycosylation for antigen-experienced vaccinees were assessed with Wilcoxon matched-pairs signed rank test. Differences between naive and antigen-experienced vaccinees two unpaired groups were assessed with Mann-Whitney U test and differences in sialylation and galactosylation for fucosylated and afucosylated anti-S IgG1 were determined by a paired t-test.The correlations between anti-S IgG1 titer and afucosylation was determined by linear regression. *P*-values < 0.05 were considered as significant. Asterisks indicate the degree of significance as follows: *, **, ***, ****: *p*-value < 0.05, 0.01, 0.001, 0.0001, respectively.

## Acknowledgements

We thank the Academic Medical Centre of the University of Amsterdam, the Sanquin Blood Supply Foundation and The Fatebenefratelli-Sacco Infectious Diseases Physicians Group. We are greatly indebted to all cohort participants for their extensive participation.

## Funding

Landsteiner foundation for Blood Transfusion Research (LSBR) grants 1721 and 1908 (GV) ZonMW COVID-19 grants 1043001 201 0021 (GV)

Deutsche Forschungsgemeinschaft (DFG, German Research Foundation) 398859914 (EH 221/10-1); 400912066 (EH 221/11-1); and 390884018 (Germany’s Excellence Strategies - EXC 2167, Precision Medicine in Chronic Inflammation (PMI)) (ME)

Federal State Schleswig-Holstein, Germany (COVID-19 Research Initiative Schleswig-Holstein; DOI4-Nr. 3) (ME)

European Union’s Horizon 2020 research and innovation program H2020-MSCA-ITN grant agreement number 721815 (TP)

Netherlands Organization for Health Research and Development ZonMw & the Amsterdam UMC Corona Research Fund (Amsterdam UMC COVID-19 S3/HCW study group)

The Netherlands Organisation for Health Research and Development ZonMW VENI grant, Grant number 09150161910033, (LAVV)

## Authors’ contributions

Conceptualization: GV, MW, ME, JDD, APJV, ECVDS, JVC, JR

Methodology: GV, MW, WW, JDD, JN, ME, JVC, TP, HJC, JR, ASL, TLJvO, JSB

Formal analysis: JVC, TP, JR, HJC, TLJvO, TS

Investigation: JVC, TP, JR, JSB, ASL, TS, MS, WW, JN, HJC, IK, TLJvO, ELM, SL, VvK, CK, HBL

Resources: FL, RV, CG, APJV, LAVV, WWH, MAS, RM, TG, MKB, JJS, Fatebenefratelli-Sacco Infectious Diseases Physicians, and the UMC COVID-19 S3/HCW study group

Data curation: JVC, TP, JR, HJC, JSB, TLJvO

Writing – Original draft: GV, MW, ME, ECVDS, JVC, TP, JR

Writing – Review and editing: GV, MW, ME, JD, ECVDS, JVC, TP, JR, HJC, CG, LAV, JSB, TS, TLJvO, MS, WWH, WW, ASL, JN, SK, FL, RV, ML, ELM, IK, SL, VvK, CK, HBL,

MDW, NVM, TR, RG, MAS, RM, APJV, Fatebenefratelli-Sacco Infectious Disease Physicians, and the UMC COVID-19 S3/HCW study group

Visualization: JVC, HJC, JSB, TP, TLJvO Supervision: GV, MW, ME, JVD

Project administration: GV, MW, ME, JVC, JVC Funding acquisition: GV, MW, ME, JVD

## Supplementary

**Table S1.**
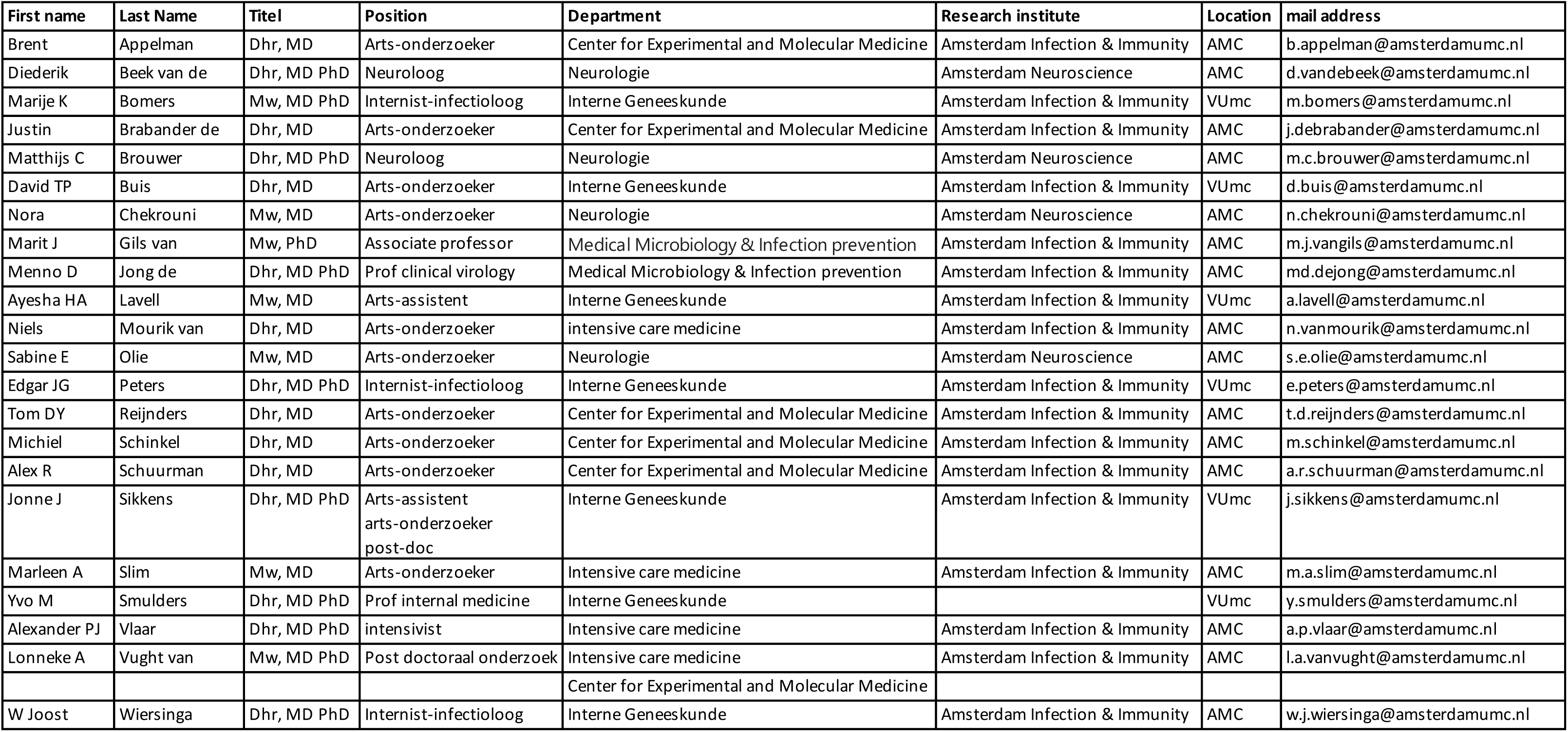
UMC COVID-19 S3/HCW study group.

**Table S2.**
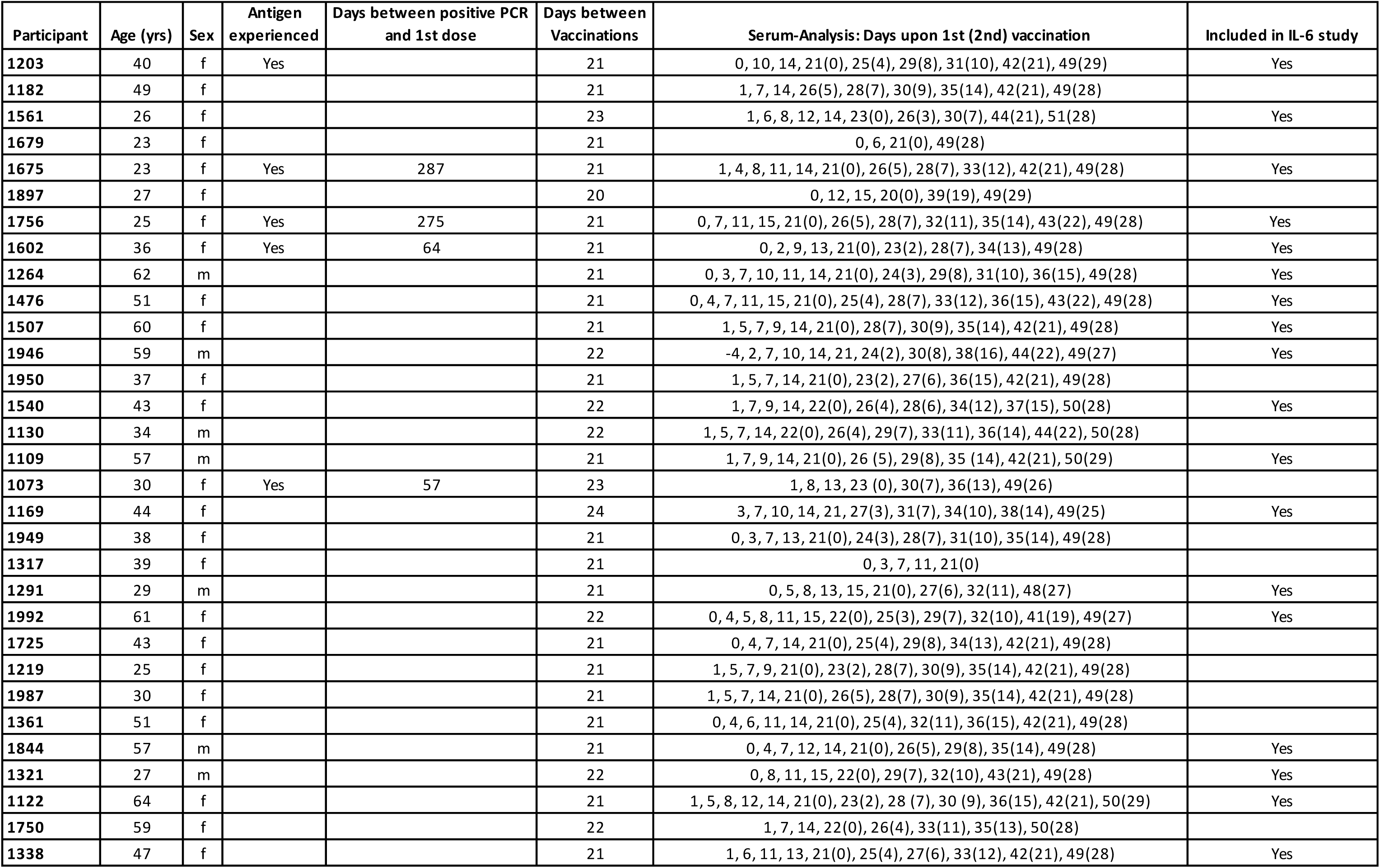
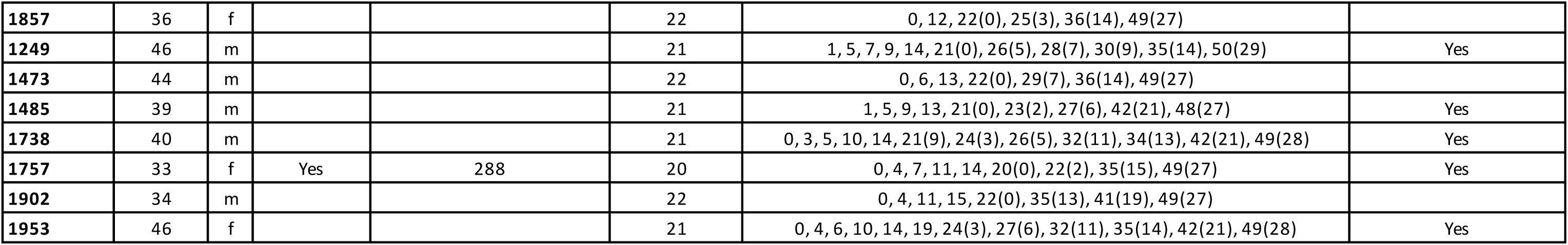
Detailed description of Cohort 1 - Amsterdam UMC cohort.

| Participant | Age (yrs) | Sex | Antigen experienced | Days between positive PCR and 1st dose | Days between Vaccinations | Serum-Analysis: Days upon 1st (2nd) vaccination | Included in IL-6 study |
| --- | --- | --- | --- | --- | --- | --- | --- |
| 1203 | 40 | f | Yes |  | 21 | 0, 10, 14, 21(0), 25(4), 29(8), 31(10), 42(21), 49(29) | Yes |
| 1182 | 49 | f |  |  | 21 | 1, 7, 14, 26(5), 28(7), 30(9), 35(14), 42(21), 49(28) |  |
| 1561 | 26 | f |  |  | 23 | 1, 6, 8, 12, 14, 23(0), 26(3), 30(7), 44(21), 51(28) | Yes |
| 1679 | 23 | f |  |  | 21 | 0, 6, 21(0), 49(28) |  |
| 1675 | 23 | f | Yes | 287 | 21 | 1, 4, 8, 11, 14, 21(0), 26(5), 28(7), 33(12), 42(21), 49(28) | Yes |
| 1897 | 27 | f |  |  | 20 | 0, 12, 15, 20(0), 39(19), 49(29) |  |
| 1756 | 25 | f | Yes | 275 | 21 | 0, 7, 11, 15, 21(0), 26(5), 28(7), 32(11), 35(14), 43(22), 49(28) | Yes |
| 1602 | 36 | f | Yes | 64 | 21 | 0, 2, 9, 13, 21(0), 23(2), 28(7), 34(13), 49(28) | Yes |
| 1264 | 62 | m |  |  | 21 | 0, 3, 7, 10, 11, 14, 21(0), 24(3), 29(8), 31(10), 36(15), 49(28) | Yes |
| 1476 | 51 | f |  |  | 21 | 0, 4, 7, 11, 15, 21(0), 25(4), 28(7), 33(12), 36(15), 43(22), 49(28) | Yes |
| 1507 | 60 | f |  |  | 21 | 1, 5, 7, 9, 14, 21(0), 28(7), 30(9), 35(14), 42(21), 49(28) | Yes |
| 1946 | 59 | m |  |  | 22 | -4, 2, 7, 10, 14, 21, 24(2), 30(8), 38(16), 44(22), 49(27) | Yes |
| 1950 | 37 | f |  |  | 21 | 1, 5, 7, 14, 21(0), 23(2), 27(6), 36(15), 42(21), 49(28) |  |
| 1540 | 43 | f |  |  | 22 | 1, 7, 9, 14, 22(0), 26(4), 28(6), 34(12), 37(15), 50(28) | Yes |
| 1130 | 34 | m |  |  | 22 | 1, 5, 7, 14, 22(0), 26(4), 29(7), 33(11), 36(14), 44(22), 50(28) |  |
| 1109 | 57 | m |  |  | 21 | 1, 7, 9, 14, 21(0), 26 (5), 29(8), 35 (14), 42(21), 50(29) | Yes |
| 1073 | 30 | f | Yes | 57 | 23 | 1, 8, 13, 23 (0), 30(7), 36(13), 49(26) |  |
| 1169 | 44 | f |  |  | 24 | 3, 7, 10, 14, 21, 27(3), 31(7), 34(10), 38(14), 49(25) | Yes |
| 1949 | 38 | f |  |  | 21 | 0, 3, 7, 13, 21(0), 24(3), 28(7), 31(10), 35(14), 49(28) |  |
| 1317 | 39 | f |  |  | 21 | 0, 3, 7, 11, 21(0) |  |
| 1291 | 29 | m |  |  | 21 | 0, 5, 8, 13, 15, 21(0), 27(6), 32(11), 48(27) | Yes |
| 1992 | 61 | f |  |  | 22 | 0, 4, 5, 8, 11, 15, 22(0), 25(3), 29(7), 32(10), 41(19), 49(27) | Yes |
| 1725 | 43 | f |  |  | 21 | 0, 4, 7, 14, 21(0), 25(4), 29(8), 34(13), 42(21), 49(28) |  |
| 1219 | 25 | f |  |  | 21 | 1, 5, 7, 9, 21(0), 23(2), 28(7), 30(9), 35(14), 42(21), 49(28) |  |
| 1987 | 30 | f |  |  | 21 | 1, 5, 7, 14, 21(0), 26(5), 28(7), 30(9), 35(14), 42(21), 49(28) |  |
| 1361 | 51 | f |  |  | 21 | 0, 4, 6, 11, 14, 21(0), 25(4), 32(11), 36(15), 42(21), 49(28) |  |
| 1844 | 57 | m |  |  | 21 | 0, 4, 7, 12, 14, 21(0), 26(5), 29(8), 35(14), 49(28) | Yes |
| 1321 | 27 | m |  |  | 22 | 0, 8, 11, 15, 22(0), 29(7), 32(10), 43(21), 49(28) | Yes |
| 1122 | 64 | f |  |  | 21 | 1, 5, 8, 12, 14, 21(0), 23(2), 28 (7), 30 (9), 36(15), 42(21), 50(29) | Yes |
| 1750 | 59 | f |  |  | 22 | 1, 7, 14, 22(0), 26(4), 33(11), 35(13), 50(28) |  |
| 1338 | 47 | f |  |  | 21 | 1, 6, 11, 13, 21(0), 25(4), 27(6), 33(12), 42(21), 49(28) | Yes |
| **1857** | 36 | f |  |  | 22 | 0, 12, 22(0), 25(3), 36(14), 49(27) |  |
| **1249** | 46 | m |  |  | 21 | 1, 5, 7, 9, 14, 21(0), 26(5), 28(7), 30(9), 35(14), 50(29) | Yes |
| **1473** | 44 | m |  |  | 22 | 0, 6, 13, 22(0), 29(7), 36(14), 49(27) |  |
| **1485** | 39 | m |  |  | 21 | 1, 5, 9, 13, 21(0), 23(2), 27(6), 42(21), 48(27) | Yes |
| **1738** | 40 | m |  |  | 21 | 0, 3, 5, 10, 14, 21(9), 24(3), 26(5), 32(11), 34(13), 42(21), 49(28) | Yes |
| **1757** | 33 | f | Yes | 288 | 20 | 0, 4, 7, 11, 14, 20(0), 22(2), 35(15), 49(27) | Yes |
| **1902** | 34 | m |  |  | 22 | 0, 4, 11, 15, 22(0), 35(13), 41(19), 49(27) |  |
| **1953** | 46 | f |  |  | 21 | 0, 4, 6, 10, 14, 19, 24(3), 27(6), 32(11), 35(14), 42(21), 49(28) | Yes |

**Table S3.** Detailed description of Cohort 2 - Fatebenefratelli-Sacco Infectious Diseases Physicians Group.

| Participant | Age (yrs) | Sex | Days between first and second dose | BMI | Cormobidities | Medications | Serum-Analysis: Days upon 1st (2nd) dose |
| --- | --- | --- | --- | --- | --- | --- | --- |
| VC-001 | 58 | M | 21 | 26 | None | None | 0, 3, 7, 11, 14, 17, 21(0), 24(3), 28(7), 32(11), 35(14), 39(18), 42(21) |
| SCM | 28 | M | 21 | 21 | None | None | 0, 4, 8, 11, 15, 18, 21(0), 25(4), 29(8), 31(10), 36(15), 39(18), 42(21) |
| TOA | 35 | M | 21 | 23 | None | None | -1, 3, 7, 10, 14, 17, 20, 24(3), 28(7), 30(9), 35(14), 38(17), 41(20) |
| ANSP | 62 | M | 21 | 24 | Hypertension<br>Hereditary spherocytosis<br>splenectomy | Olmesartan<br>Esomeprazole | 0, 4, 8, 11, 15, 18, 22(1), 25(4), 29(8), 32(11), 35(14), 43(22), 45(24) |
| COA | 28 | F | 21 | 21 | None | None | -1, 3, 6, 10, 13, 17, 20, 24(3), 27(6), 31(10), 34(13), 38(17), 41(20) |
| BAC | 29 | F | 21 | 19 | None | None | -1, 3, 6, 10, 13, 17, 20, 25(4), 27(6), 31(10), 34(13), 38(17), 41(20) |
| LUAN | 59 | F | 21 | 31 | Hypertension | Ramipril | 0, 4, 8, 11, 15, 18, 22(1), 25(4), 32(11), 34(13), 36(15), 39(18), 43(22) |
| MIL | 53 | F | 21 | 19 | Breast Cancer | Aromasin | -1, 2, 6, 9, 13, 16, 20, 23(2), 27(6), 30(9), 34(13), 37(16), 41(20) |
| BEGI | 55 | F | 21 | 29 | None | None | 0, 3, 7, 10, 14, 17, 21, 24(3), 28(7), 31(10), 35(14), 38(17), 42(21) |

**Table S4.**
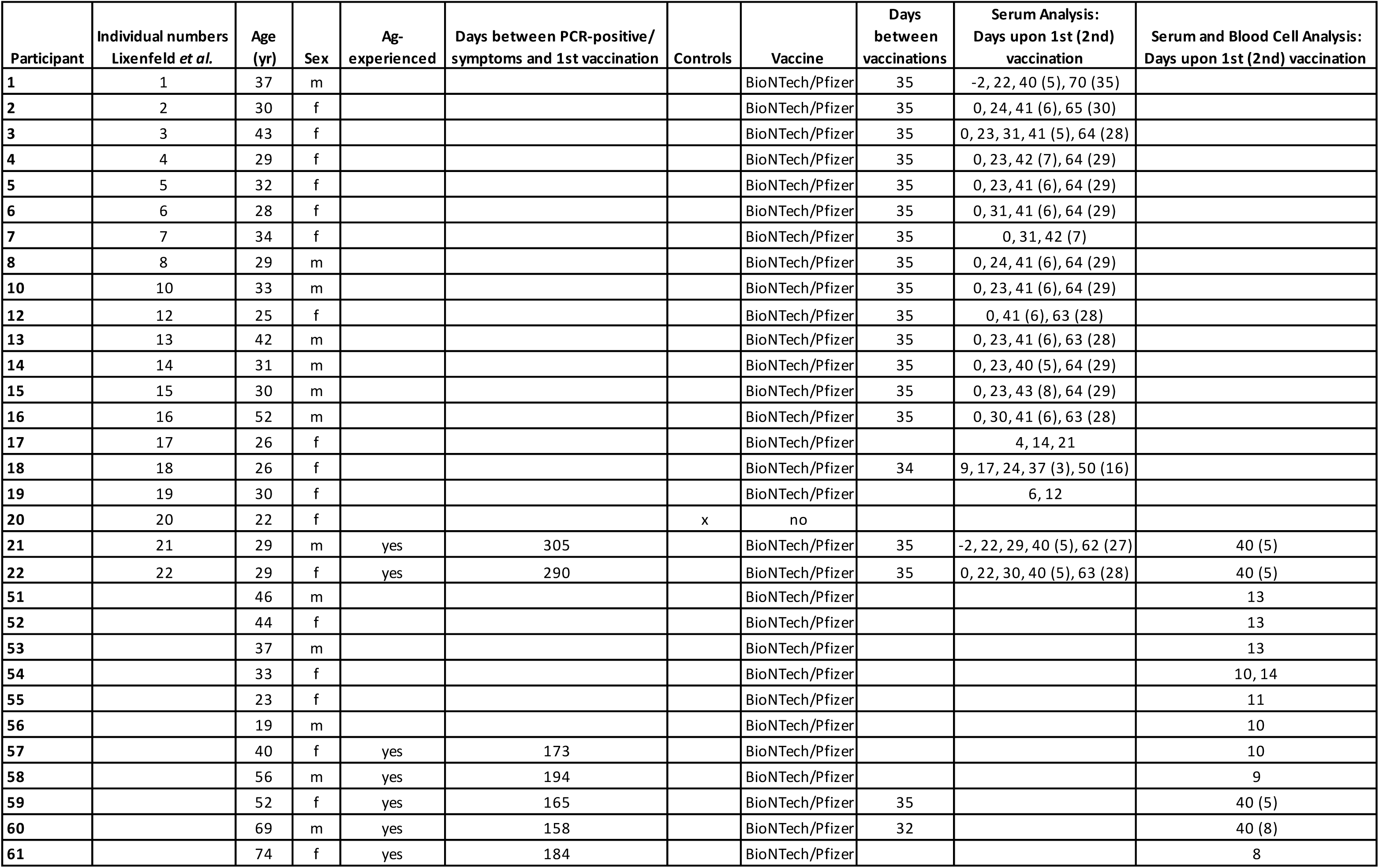
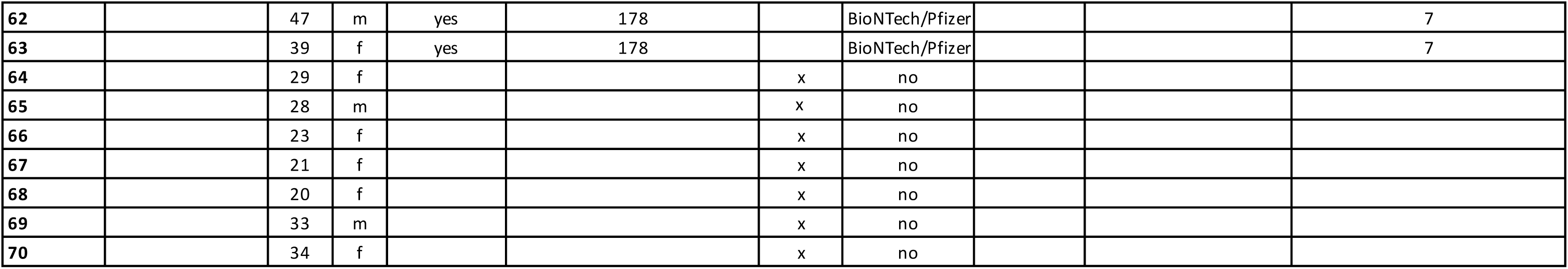
Detailed description of Cohort 3 - Luebeck cohort.

**Table S5.** Detailed description of Cohort 4 - Convalescent plasma donors.

| **Participant** | **Age (yrs)** | **Date of pre-vaccination sample** | **Date of post-vaccination sample** |
| --- | --- | --- | --- |
| 70826 | 48 | 31/12/2020 | 15/02/2021 |
| 511828 | 56 | 21/12/2020 | 22/02/2021 |
| 553113 | 44 | 31/12/2020 | 10/02/2021 |
| 616886 | 49 | 15/01/2021 | 24/02/2021 |
| 632961 | 62 | 30/12/2020 | 24/02/2021 |
| 659201 | 41 | 03/02/2021 | 24/02/2021 |
| 1086369 | 53 | 14/01/2021 | 22/02/2021 |
| 1217364 | 49 | 21/01/2021 | 11/02/2021 |
| 1218801 | 59 | 18/12/2020 | 15/02/2021 |
| 1339188 | 47 | 05/01/2021 | 22/02/2021 |
| 1832436 | 30 | 04/01/2021 | 24/02/2021 |
| 1878273 | 28 | 18/01/2021 | 16/02/2021 |
| 1937124 | 27 | 06/01/2021 | 25/02/2021 |
| 1972705 | 51 | 12/01/2021 | 03/02/2021 |
| 2177185 | 25 | 19/01/2021 | 22/02/2021 |
| 2198851 | 60 | 06/01/2021 | 10/02/2021 |
| 2394658 | 35 | 05/11/2020 | 02/02/2021 |
| 2402217 | 45 | 21/01/2021 | 11/02/2021 |
| 2408917 | 57 | 13/01/2021 | 03/02/2021 |
| 2411367 | 59 | 05/01/2021 | 26/01/2021 |
| 2412091 | 19 | 07/01/2021 | 28/01/2021 |
| 2427249 | 25 | 07/01/2021 | 26/01/2021 |

**Table S6.**
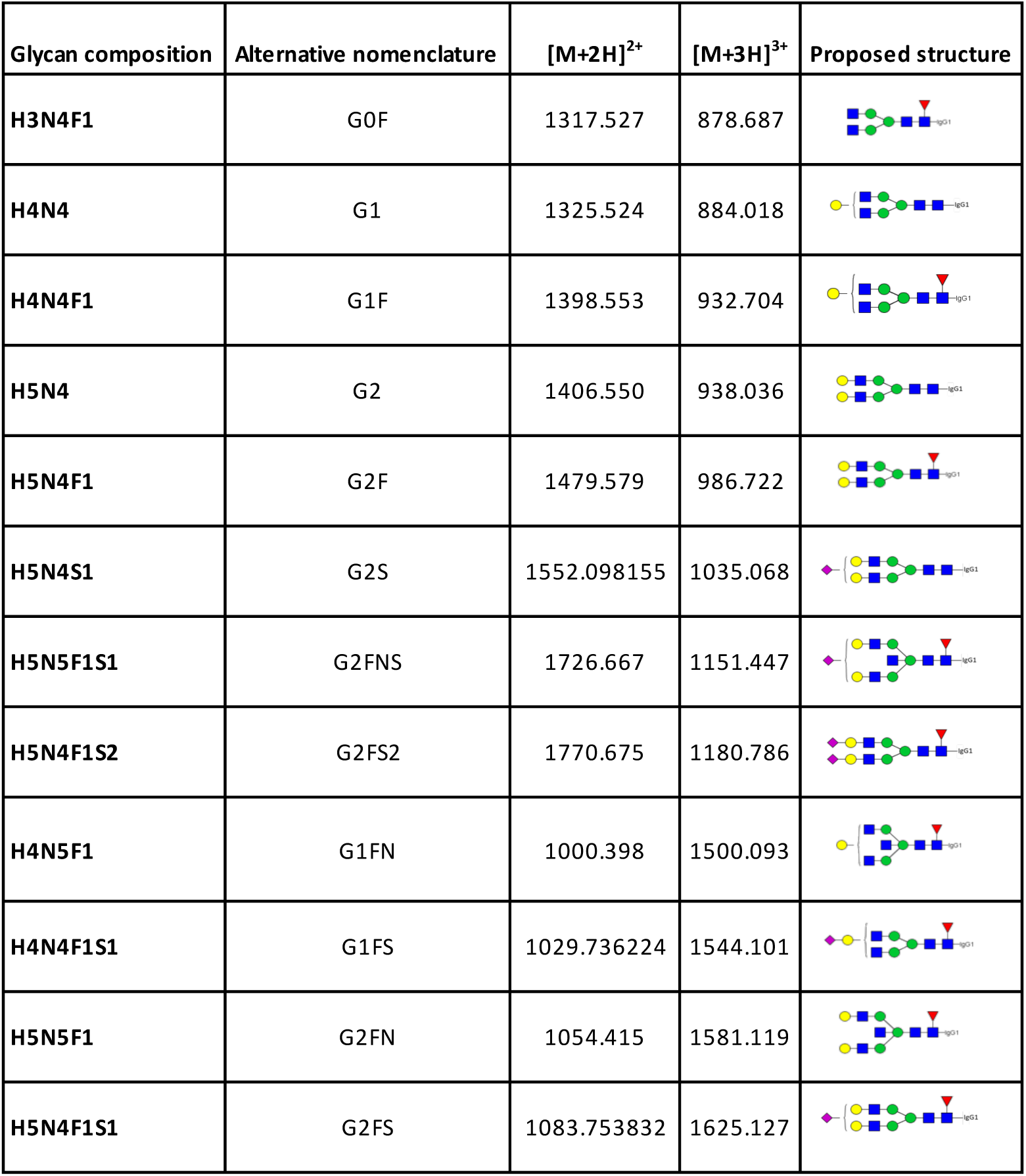
IgG1 glycopeptides included in the final analyte list.

| Glycan composition | Alternative nomenclature | [M+2H]<sup>2+</sup> | [M+3H]<sup>3+</sup> | Proposed structure |
| --- | --- | --- | --- | --- |
| H3N4F1 | G0F | 1317.527 | 878.687 |  |
| H4N4 | G1 | 1325.524 | 884.018 |  |
| H4N4F1 | G1F | 1398.553 | 932.704 |  |
| H5N4 | G2 | 1406.550 | 938.036 |  |
| H5N4F1 | G2F | 1479.579 | 986.722 |  |
| H5N4S1 | G2S | 1552.098155 | 1035.068 |  |
| H5N5F1S1 | G2FNS | 1726.667 | 1151.447 |  |
| H5N4F1S2 | G2FS2 | 1770.675 | 1180.786 |  |
| H4N5F1 | G1FN | 1000.398 | 1500.093 |  |
| H4N4F1S1 | G1FS | 1029.736224 | 1544.101 |  |
| H5N5F1 | G2FN | 1054.415 | 1581.119 |  |
| H5N4F1S1 | G2FS | 1083.753832 | 1625.127 |  |

**Table S7.** Description and calculation of IgG1 glycosylation traits. H: hexose, N: N-acetylhexosamine, F: fucose, S: N-acetylneuraminic (sialic) acid.

|  | Description | Formula |
| --- | --- | --- |
| IgG1 bisection | *N*-glycans carrying a bisected *N*-acetylglucoseamine | ( H5N5F1S1 + H4N5F1 + H5N5F1 ) / sum of all IgG1 glycopeptides |
| IgG1 galactosylation | *N*-glycans carrying galactose(s) | ( 1/2 * ( H4N4 + H4N4F1 + H4N5F1 + H4N4F1S1 ) + 2/2 * ( H5N4 + H5N4F1 + H5N4S1 + H5N5F1S1 + H5N4F1S2 + H5N5F1 + H5N4F1S1 ) ) / sum of all glycopeptides |
| IgG1 sialylation | *N*-glycans carrying *N*-acetylneuraminic (sialic) acid(s) | ( 1/2 * (H5N4S1 + H5N5F1S1 + H4N4F1S1 ) + 2/2 * H5N4F1S2 ) / sum of all IgG1 glycopeptides |
| IgG1 fucosylation | *N*-glycans carrying a core fucose | ( H3N4F1 + H4N4F1 + H5N4F1 + H5N5F1S1 + H5N4F1S2 + H4N5F1 + H4N4F1S1 + H5N5F1S1 + H5N4F1S1 ) / sum of all IgG1 glycopeptides |

**Figure S1.**
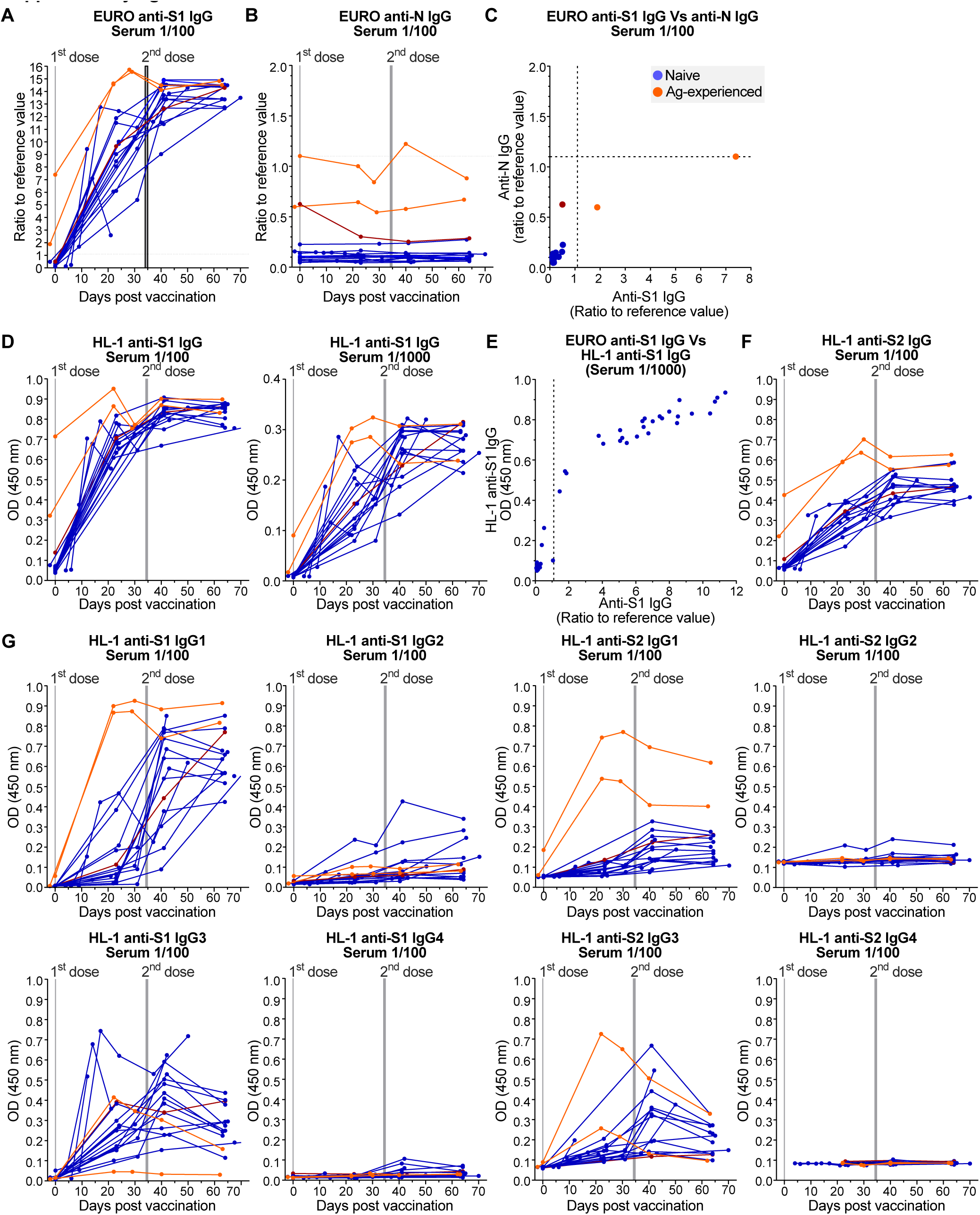
Anti-S and -N serum IgG Ab levels of cohort 3. **(A-B)** Sera of naive (blue) and antigen-experienced (yellow) individuals from a subset of cohort 3 (see Table S4) were analyzed by EUROIMMUN (EURO) anti-SARS-CoV-2-Spike1 (S1) and –Nucleocapsid (N) IgG ELISA. **(C)** Correlation between EURO the anti-S1 and –N IgG levels before vaccination. Dotted lines are reference values as determined by the company. **(D)** HL-1 anti-S1 IgG ELISA. **(E)** Correlation between the EUROIMMUN and HL-1 anti-S1 IgG ELISA data. **(F)** HL-1 anti-S2 IgG ELISA. **(G)** HL-1 anti-S1 and -S2 IgG1-4 ELISA. The data of the naive-considered individual that showed enhanced anti-N IgG levels before vaccination were marked in dark red in all graphs.

**Figure S2.**
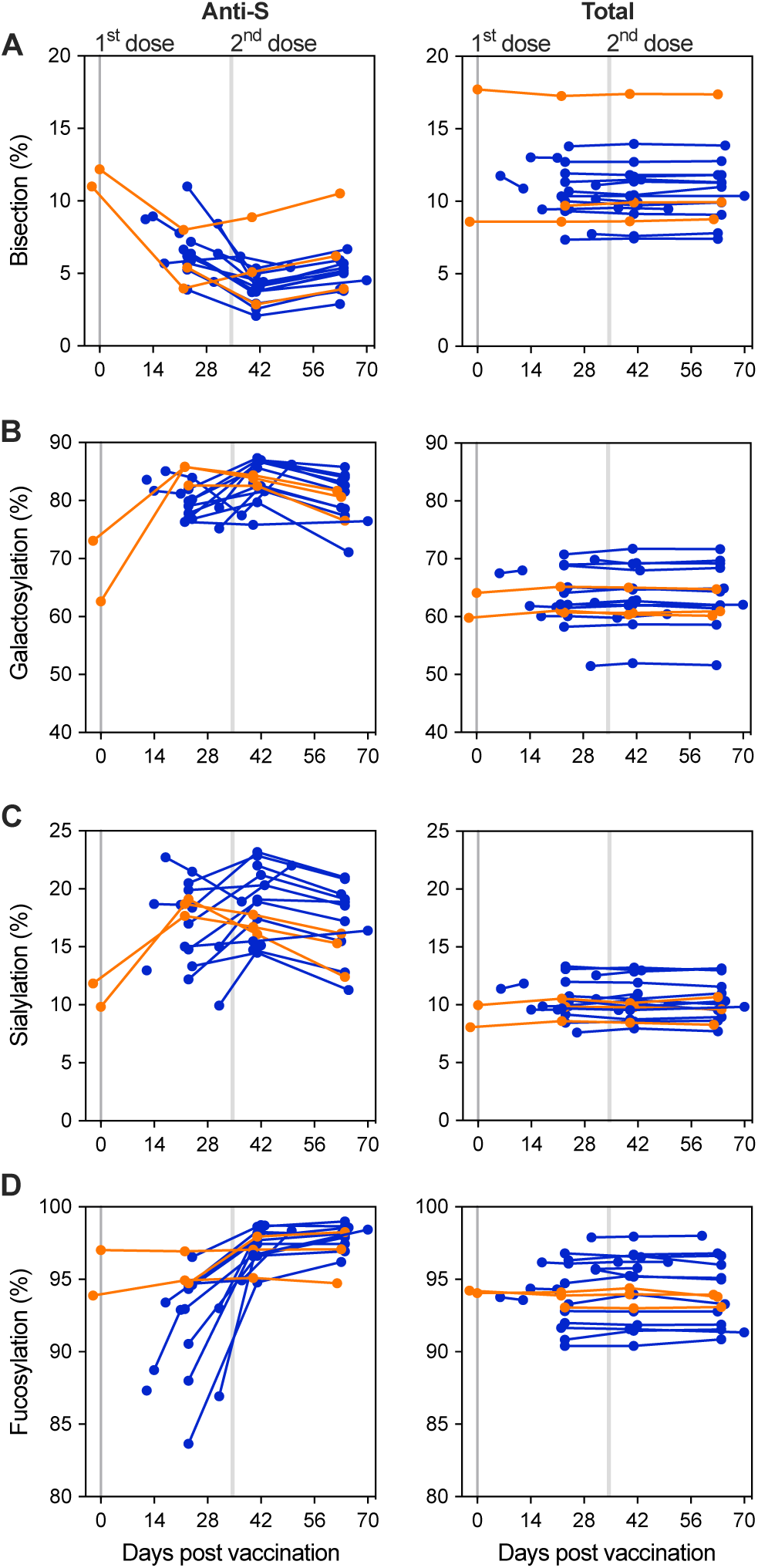
IgG1 glycoprofiling data of cohort 3. Longitudinal anti-spike (S) (left) and total (right) IgG1-Fc glycosylation for a subset of cohort 3 (See Table S4) with **(A)** bisection, **(B)** galactosylation, **(C)** sialylation, and **(D)** fucosylation of naive (blue, circle) and antigen-experienced (yellow, triangle) vaccinated participants.

**Figure S3.**
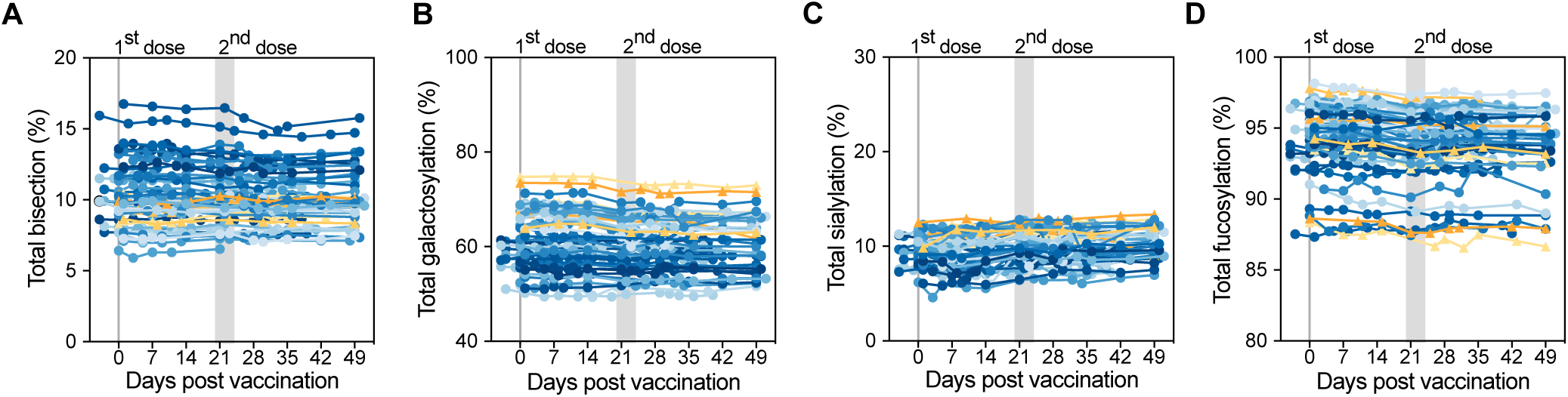
Total IgG1 Fc glycoprofiling data of cohorts 1 and 2. Longitudinal total IgG1-Fc glycan traits for cohort 1 (n=39) and 2 (n=9) with **(A)** bisection, **(B)** galactosylation, **(C)** sialylation, and **(D)** fucosylation of naive (blue) and antigen-experienced (yellow) vaccinated participants.

**Figure S4.**
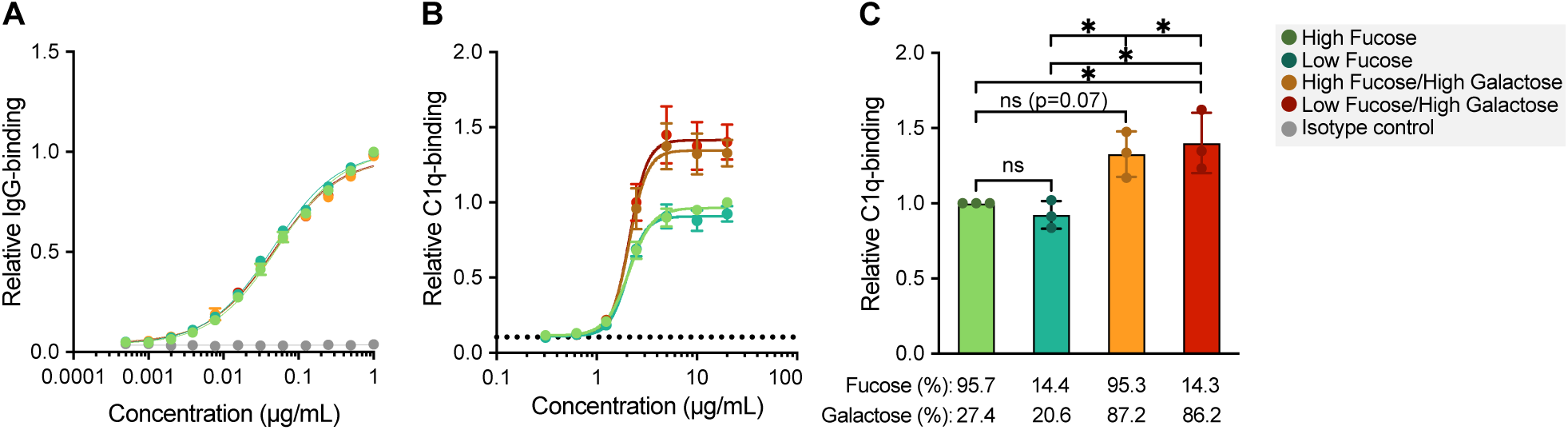
Complement activation of glycoengineered anti-S COVA1-18 mAb. Glycoengineered anti-S COVA1-18 was produced with high/low fucosylation in combination with normal and increased galactosylation. **(A)** Binding to the receptor-binding domain (RBD) of the spike (S) protein. **(B)** relative binding of C1q and **(C)** relative levels of C1q binding presented as a relative value to the maximum response of the unmodified WT anti-S COVA1-18 (2C1) hIgG1 mAb. Levels were determined by ELISA (n=3) and curve fitting was performed using nonlinear regression dose-response curves with log(agonist) versus response–variable slope (four parameters) (* *p*-value < 0.05).

**Figure S5.**
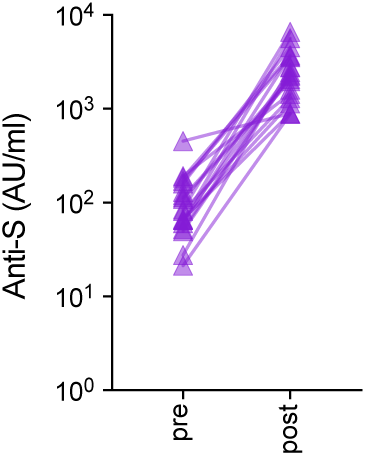
Anti-S IgG titers of antigen-experienced vaccinated blood donors. Anti-S IgG titers of antigen-experienced vaccinated blood donors (cohort 4, n=22) before (pre) and after (post) vaccination.

**Figure S6.**
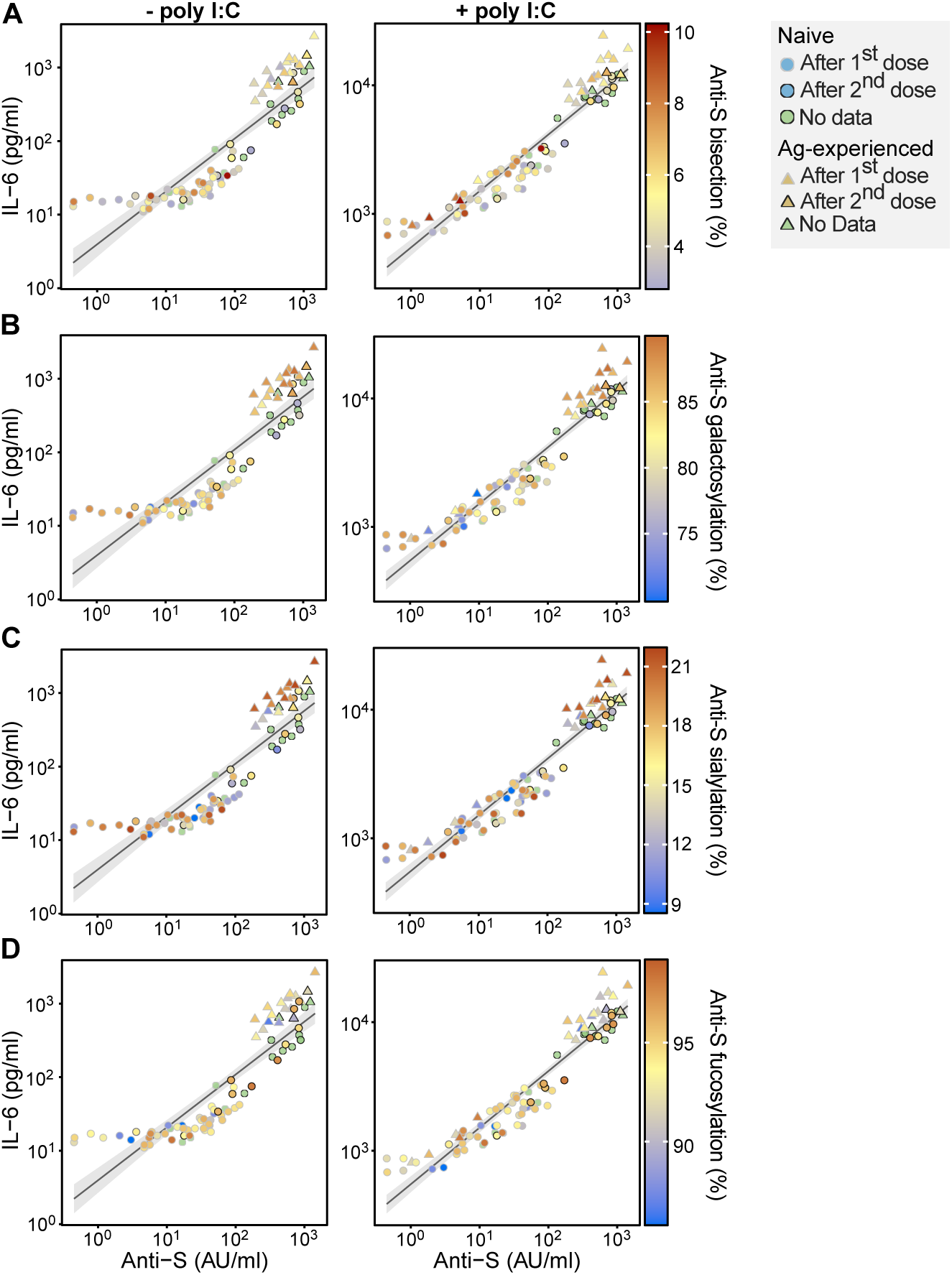
Anti-S Fc glycosylation traits have a minor influenceon IL-6 production upon macrophage stimulation. IL- 6 responses of macrophages stimulated with spike (S) protein and patient in the absence (left) or presence (right) of poly(I:C) correlated with anti-S titer. All data represent different time points of a subgroup of cohort 1 (n=23) with a gradient of anti-S IgG1-Fc **(A)** bisection, **(B)** galactosylation, **(C)** sialylation, and **(D)** fucosylation

**Figure S7.**
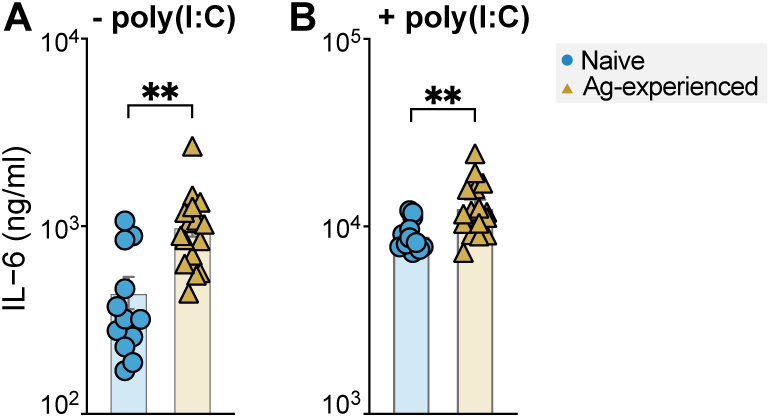
IL-6 production of naive and antigen-experienced vaccinees when titer is comparable. Comparison by unpaired *t*-test between the IL-6 responses of macrophages stimulated with spike (S) protein and naive (blue) or antigen-experienced (yellow) vaccinee sera when titers are comparable (>200 AU/ml) in the **(A)** absence or **(B)** presence of poly(I:C). All data represent a subgroup of cohort 1 (n=23, see Table S2).

**Figure S8.**
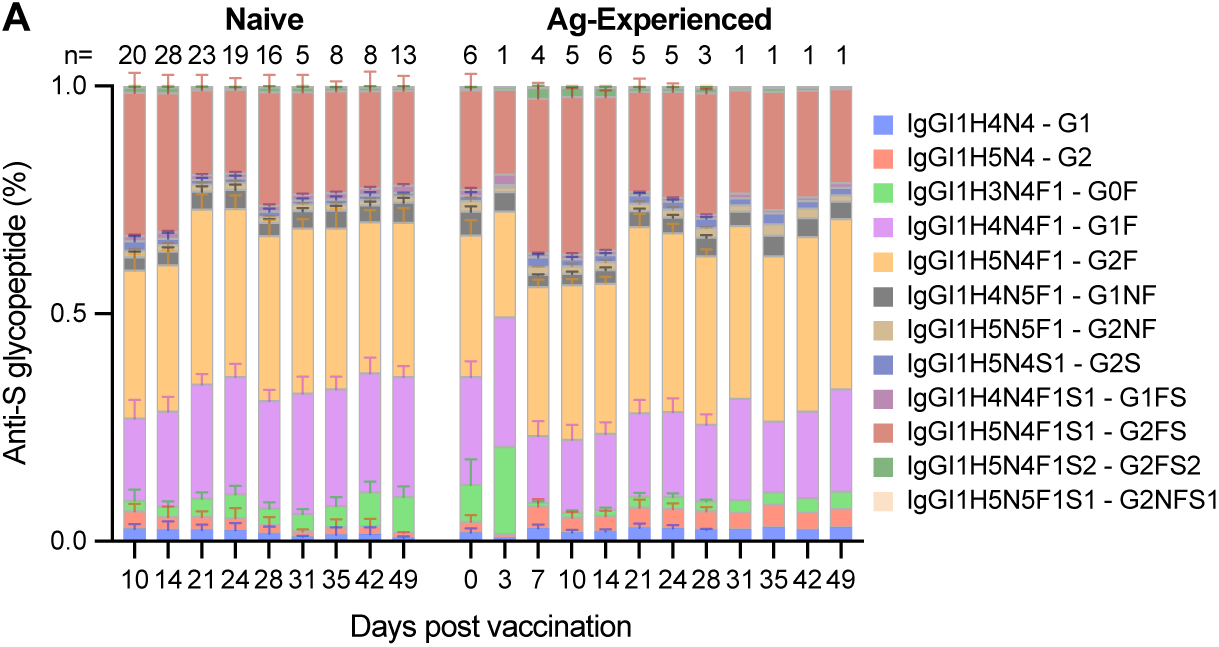
Anti-S IgG1 Fc glycopeptide composition. Anti-spike (S) IgG1-Fc glycopeptides composition for naive (left, cohort 1 (n=33) and 2 (n=9)) and antigen-experienced (right, cohort 1 (n=6) and 2 (n=0)) over time. H: hexose; N: *N*-acetylhexosamine; F: fucose; S: *N*-acetylneuraminic (sialic) acid.

**Figure S9.**
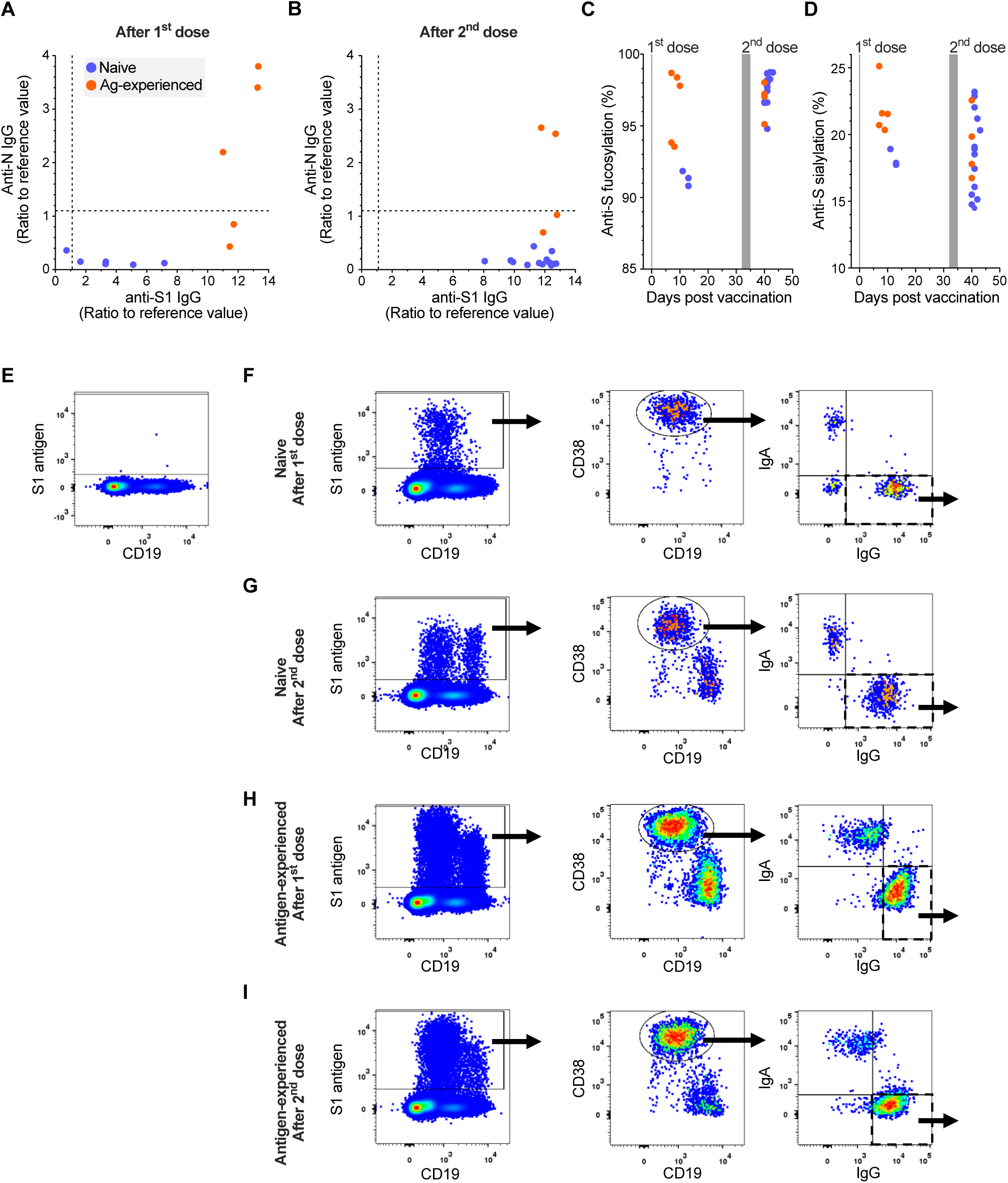
Characterization of naive and antigen-experienced individuals described in Fig. 4C-N. Serum IgG and blood cells of naive (blue) and antigen-experienced (Ag. exp.; yellow) individuals were analyzed 7-14 days after the first (naive: n=6; Ag.-exp.: n=5) or 5-8 days after the second (naive: n=15; Ag.-exp.: n=4) dose by ELISA and flow cytometry. **(A, B)** Correlation of EUROIMMUN anti-SARS-CoV-2-S1 and -N IgG values from samples after the **(A)** first or **(B)** second dose. **(C)** Anti-S IgG1 fucosylation and **(D)** anti-S1 IgG1 sialylation of all samples. **(E-I)** Blood cells were gated on single, living lymphocytes. Example gating strategy with samples of a naive individual **(E)** unimmunized, **(F)** after the 1^st^ dose and **(G)** after the 2^nd^ dose and an antigen-experienced individual after **(H)** the 1^st^ dose and **(I)** the 2^nd^ dose with BNT162b2. S1-reactive B cells were gated and further gated for CD19^int^ CD38^high(+)^ PCsto analyze IgG^+^ PCs.

**Figure S10.**
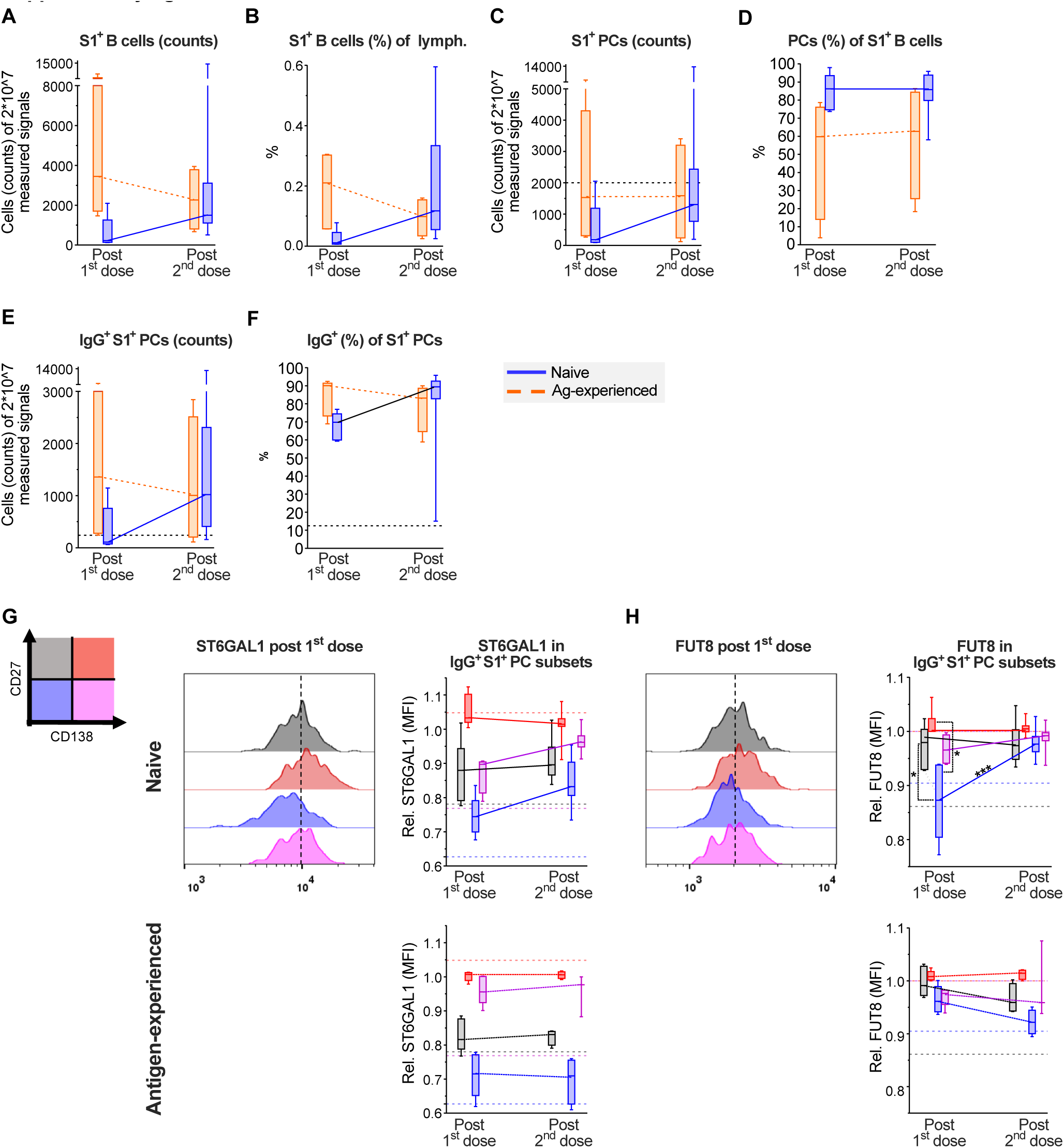
Counts, frequencies and glycosyltransferase expression of IgG^+^ S1-reactive PC subsets from naive and antigen-experienced individuals described in Fig. 4C-N and S8. (A-F) Counts and frequencies of S1-reactive B cells, S1-reactive CD38^+^ PCs and IgG^+^ S1-reactive CD38^+^ PCs from naive (blue) and antigen-experienced (yellow) individuals. (G, H) IgG^+^ S1-reactive CD38^+^ PCs were subdivided in CD27^+^ CD138^-^ (grey), CD27^+^CD138^+^ (red), CD27^low^CD138^-^ (blue) and CD27^low^CD138^+^ (pink) subsets and analyzed for (G) ST6GAL1 or (H) FUT8 expression. Example overlay histograms and the relative glycosyltransferase expression (median (MFI) of all samples are shown. The median (MFI) of ST6GAL1 or FUT8 expression in CD138^+^ IgG+ S1-reactive PCs of each sample was set to 1 for inter-assay comparison. The fine dotted horizontal lines indicate corresponding values of total IgG+ PC subsets from untreated healthy controls (Fig. S10).

**Figure S11.**
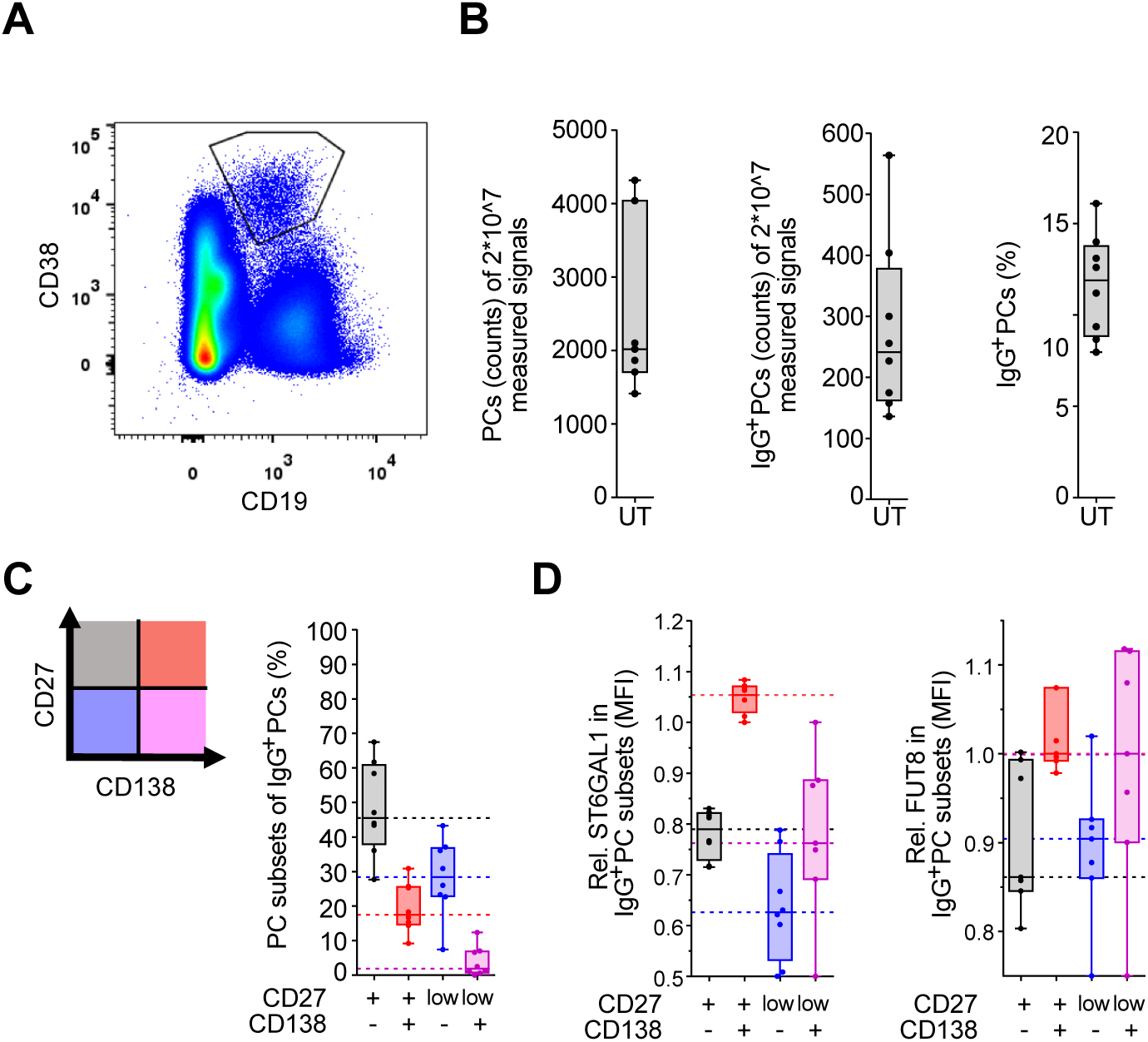
Counts, frequencies, and glycosyltransferase expression of PCs from untreated healthy individuals. Blood cells of untreated (UT) healthy individuals (n=8) were analyzed by flow cytometry. Blood cells were gated on single, living lymphocytes. **(A)** Example gating strategy of CD19^int^ CD38^high(+)^ PCs. **(B, C)** Counts and frequencies of PCs, IgG^+^ PCs and CD27^+^CD138^-^ (grey), CD27^+^CD138^+^ (red), CD27^low^CD138^-^ (blue) and CD27^low^CD138^+^ (pink) IgG^+^ PC subsets gated as described in Fig. 4, S8 and S9. **(D)** Relative ST6GAL1 or FUT8 expression median (MFI) in the four IgG^+^ PC subsets. The median (MFI) of ST6GAL1 or FUT8 expression in CD138^+^ IgG+ PCs of each sample was set to 1 for inter-assay comparison.

**Figure S12.**
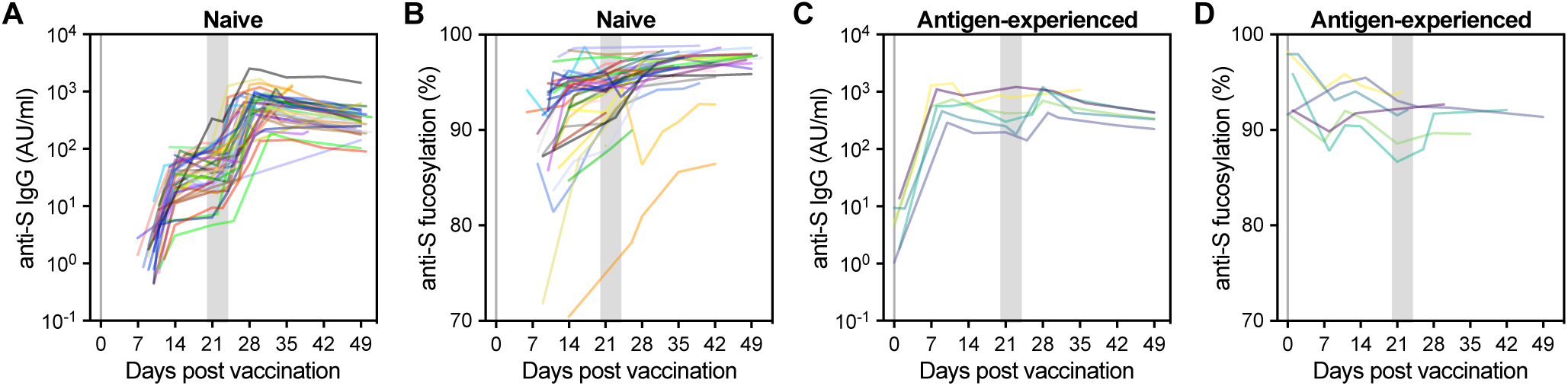
Anti-S IgG titer and IgG1-Fc glycosylation per individual. Anti-S IgG **(A, C)** titer and **(B, D)** IgG1-Fc fucosylation shown for SARS-CoV-2 **(A-B)** naive and **(C-D)** antigen-experienced vaccinees over time.

**Figure S13.**
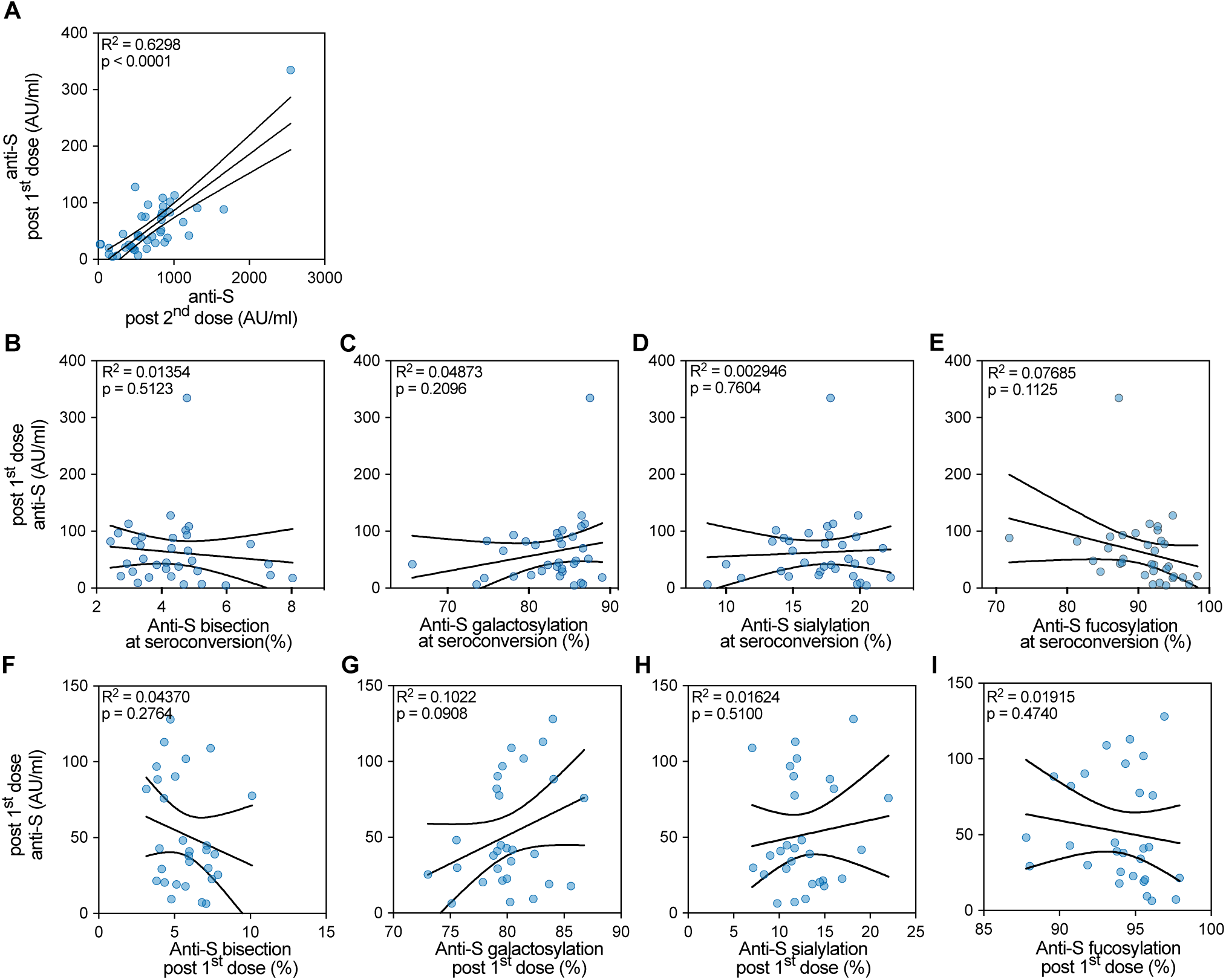
Anti-Spike IgG titer after the first and second dose correlate. **(A)** Correlation by linear regression between anti-Spike (S) titer from cohort 1 (n=39) and 2 (=8), when timing of sampling matched selection criteria, after the 1^st^ dose (on the day of the 2^nd^ dose up until three days prior) and after the 2^nd^ dose (highest titer up to 14 days after the 2^nd^ dose). **(B-I)** Correlation by linear regression between anti-Spike (S) IgG1-Fc glycoprofiling (left: bisection, middle left: galactosylation, middle right: sialylation, and right: fucosylation) and titer for naive vaccines from cohort 1 (n=33) and 2 (n=9) when the timing of sampling matched selection criteria. **(B-E)** Correlation of anti-S IgG1-Fc glycosylation upon seroconversion and **(F-I)** after the first dose (on the day of the 2^nd^ dose up until three days prior) with titer after the first dose (on the day of the 2^nd^ dose up until three days prior).

